# Dynamic regulation of Sec24C by phosphorylation and O-GlcNAcylation during cell cycle progression

**DOI:** 10.1101/2025.02.12.638000

**Authors:** George R. Georgiou, Tetsuya Hirata, Erik Soderblom, Rose Homoelle, Jakob Maiwald, Michael Boyce

**Affiliations:** Department of Biochemistry, Duke University School of Medicine, Durham, North Carolina, USA; Duke Proteomics and Metabolomics Core Facility, Center for Genomic and Computational Biology, Duke University, Durham, North Carolina, USA; Haverford College, Haverford, Pennsylvania, USA

**Keywords:** cell cycle, mitosis, COPII, intracellular trafficking, O-GlcNAcylation, phosphorylation

## Abstract

During mitosis, eukaryotic cells cease anterograde trafficking from the endoplasmic reticulum (ER) towards the Golgi. This cessation corresponds with the dispersal of the COPII transport protein, Sec24C, from juxtanuclear ER exit sites (ERES) into a diffusely cytosolic pool. Redistribution of Sec24 paralogs and other core COPII proteins may underlie the mitotic pause in secretion and may be required for the equal inheritance of endomembrane organelles and machinery by both daughter cells. Therefore, it is important to understand the mechanisms governing the mitotic relocalization of COPII components. Here, we explore the role of post-translational modifications (PTMs) of the model COPII protein Sec24C in this phenotypic switch during mitosis. In interphase, Sec24C is modified by O-linked β-*N*-acetylglucosamine (O-GlcNAc), and we show that this glycan is rapidly removed upon mitotic entry, influencing the timing of Sec24C dispersal. Additionally, we identify novel, cell cycle phase-enriched phosphorylation events on Sec24C, including phosphosites that regulate the stability and localization of the protein, providing the first systematic characterization of dynamic PTMs on any Sec24 protein. Together, our data support the hypothesis that phosphorylation and glycosylation of Sec24C act in concert to induce rapid dispersal upon mitotic entry and may promote equal partitioning of the endomembrane system to daughter cells after division.

## Introduction

The endomembrane system directs about a third of all eukaryotic proteins to their final destinations **(1–4)**. In particular, the endoplasmic reticulum (ER) selectively packages and traffics proteins and lipids to the Golgi, a process crucial for homeostasis. A multi-subunit complex, coat protein complex II (COPII), performs this trafficking **(3, 5–8)**. COPII self-organizes at specialized foci on the ER, known as ER exit sites (ERES), interacts with lumenal and membrane-bound secretory pathway cargoes, and assembles into carriers for anterograde trafficking **(3, 5–8)**. In *in vitro* budding assays, COPII carrier assembly requires five proteins: Sar1, Sec23, Sec24, Sec13, and Sec31 **(3, 5–8)**. *In vivo*, Sec12 and Sec16 are also required, and several other accessory proteins, such as cTAGE5 and TANGO1, may be necessary, depending on specific tissue and transport demands **(9–13)**.

Decades of research have elucidated much detail on the structures of COPII subunits, their biochemical interactions with one another and with some cargoes, and their assembly at ERES **(8, 14, 15)**. COPII carrier assembly begins at ERES, ribosome-free regions of rough ER marked by Sec16 **(3, 5–8)**. Sec16 is associated with Sec12, a guanine nucleotide exchange factor for Sar1, which is a soluble cytoplasmic protein when GDP-bound **(3, 5–8)**. Sec12 facilitates the exchange of GDP for GTP on Sar1, inducing an N-terminal alpha helix of Sar1-GTP to embed in the ER membrane **(3, 5–8)**. Sar1-GTP recruits Sec23-Sec24 heterodimers from the cytosol, which bind Sar1 and adjacent heterodimers, causing the ER membrane to bulge into the pre-budding complex **(3, 5–8)**. Then, through interaction with the Sec23-Sec24 inner coat, Sec13-Sec31 heterotetramers are recruited to form the COPII outer coat to complete assembly and scission of the mature COPII carrier **(3, 5–8)**.

Sec24 is the cargo-selecting subunit of COPII, acting through direct interactions with transmembrane client proteins and indirect interactions with lumenal cargoes via transmembrane adaptor proteins **(16–19)**. In many eukaryotes, Sec24 exists as multiple paralogs with distinct tissue-specific functions. The human paralogs, Sec24A-D, are differentially expressed across tissues and may interact with different repertoires of cargoes **(20–25)**. Sec24D, for example, is known to play a significant role in the transport of collagen. Indeed, in human patients, *sec24d* mutations cause Cole-Carpenter Syndrome 2 and a subtype of osteogenesis imperfecta, both severe skeletal disorders associated with the improper collagen trafficking **(26–29)**.

The existence of multiple Sec24 paralogs may provide one way to fine-tune COPII transport to specific cellular needs, but the molecular mechanisms and downstream consequences of this form of regulation remain enigmatic. Indeed, despite significant progress, key aspects of COPII trafficking remain unclear, such as the mechanism of subunit recruitment from the cytosol to ERES, the regulation of cargo selection and transport kinetics in response to cellular and tissue needs, the accommodation and transport of large cargoes, the mechanism ensuring unidirectional movement of COPII carriers towards the Golgi, and the spatiotemporal regulation of uncoating **(3, 5–8)**. As the cargo-selecting subunit of COPII and an integral component of the inner coat, Sec24 is likely central to several of these unanswered questions. New approaches are needed to dissect the regulation and function of individual Sec24 paralogs.

In recent years, discoveries have highlighted the importance of post-translational modifications (PTMs) in regulating COPII transport **(30)**. For example, in yeast, the Golgi-localized Hrr25p kinase (an ortholog of human CK1δ) phosphorylates Sec23/Sec24, triggering COPII uncoating from ER membranes and carrier fusion with the Golgi **(31)**. We and others have also shown that COPII trafficking is regulated by O-GlcNAcylation, the reversible addition of O-linked β-*N*-acetylglucosamine to serine or threonine residues of cytosolic and nuclear proteins **(30, 32–34)**. O-GlcNAc is added and removed from substrates by O-GlcNAc transferase (OGT) and O-GlcNAcase (OGA), respectively **(35–38)**. O-GlcNAc sites have been identified on thousands of proteins, including COPII subunits, with diverse and significant functional consequences **(30, 32–34, 39–42)**. Previous work in our lab, for example, found that O-GlcNAcylation of Sec23A (one of two human paralogs of Sec23) is required for endogenous collagen transport in human chondrosarcoma cells and zebrafish models **(33)**. Other work showed that O-GlcNAcylation of Sec31A (one of two human paralogs of Sec31) is important for its interaction with an essential binding partner, Sec13 **(34)**.

We have previously identified multiple PTM sites on human Sec24 proteins, but the functional implications of these observations remain largely unexplored **(33, 34)**. Interestingly, a prior study suggested that O-GlcNAcylation and phosphorylation of Sec24C together regulate COPII localization and function in a cell cycle phase-dependent manner **(43)**. COPII has long been known to respond to cell cycle cues. Studies have shown that COPII trafficking decreases significantly upon mitotic entry, and this coincides with a dramatic dispersal of COPII subunits into the cytosol **(44–46)**. PTMs may play a general role in coordinating this process, as the COPII accessory protein TANGO1 is reportedly phosphorylated in mitosis to promote ERES dissolution **(47)**. Taken together, these observations suggest that PTMs on Sec24C and other proteins may mediate the mitotic partitioning of essential endomembrane machinery to ensure equal inheritance by the daughter cells. However, despite its potential biological significance, this model has not been tested systematically.

Here, we focus on Sec24C to dissect the role of O-GlcNAcylation and phosphorylation on a critical COPII subunit during cell cycle progression. We show that Sec24C is modified by both site-specific phosphorylation and O-GlcNAcylation, with the level of each changing in a cell cycle phase-dependent manner. Furthermore, we show that the timing of mitotic dispersal of Sec24C is influenced by its glycosylation. Lastly, we identify novel Sec24C glyco- and phosphosites and provide evidence that phosphorylation at specific residues regulates Sec24C stability and localization. Our results afford new insight into the biological significance of PTMs in the fundamental control of COPII trafficking and potential paralog-specific regulation of Sec24C. More generally, our findings support the hypothesis that PTMs of trafficking proteins facilitate the inheritance of the endomembrane system after cell division **(47–58)**.

## Results

### Sec24C O-GlcNAcylation and phosphorylation change with cell cycle phase

To systematically characterize the function of Sec24C PTMs in COPII regulation, we developed a human cell model by epitope-tagging the endogenous *sec24c* locus. This strategy avoids major pitfalls of transgenesis, such as supraphysiological expression levels and non-native promoters. We used established CRISPR-based methods **(59)** to introduce a myc-6xHis tag in exon 1 of the endogenous *sec24c* gene of HeLa cells, a well-studied model system for cell cycle dynamics and regulation **(60, 61)** (Fig. S1A). Immunoblot (IB), immunoprecipitation (IP), and immunofluorescence (IF) assays demonstrated that a single cell-derived clone, dubbed “MH-Sec24C,” expressed tagged Sec24C that recapitulates the native protein-protein interactions and subcellular localization of untagged, endogenous protein (Fig. S1A-C).

First, we investigated whether Sec24C is O-GlcNAcylated in interphase and phosphorylated in mitosis, as reported previously **(43)**. MH-Sec24C HeLa cells were synchronized by double-thymidine block (DTB) (Fig. 1A) **(60)**, and synchronization was confirmed by IB for relevant markers (cyclin B1, cyclin E1, and phospho-histone H3 (Ser10) (pH3-Ser10)) (Fig. 1B) and by flow cytometry measurements of DNA content (Fig. S2A). Cell cycle phase-specific Sec24C glycosylation was quantified by IP and fluorescent IB using two distinct anti-O-GlcNAc monoclonal antibodies, clones 9D1 and 18B10 (Fig. 1B). Sec24C O-GlcNAc levels rose in G2 before falling during mitosis and then rising again upon mitotic exit and progression into G1 (Fig. 1C-D). IP-IB revealed an upper Sec24C band in mitotic but not interphase samples (Fig. 1B), consistent with a prior observation by Dudognon *et al*. that was hypothesized to result from mitotic phosphorylation of Sec24C **(43)**. Sec24C protein levels did not vary significantly across cell cycle phases (Fig. S2B).

**Figure 1.**
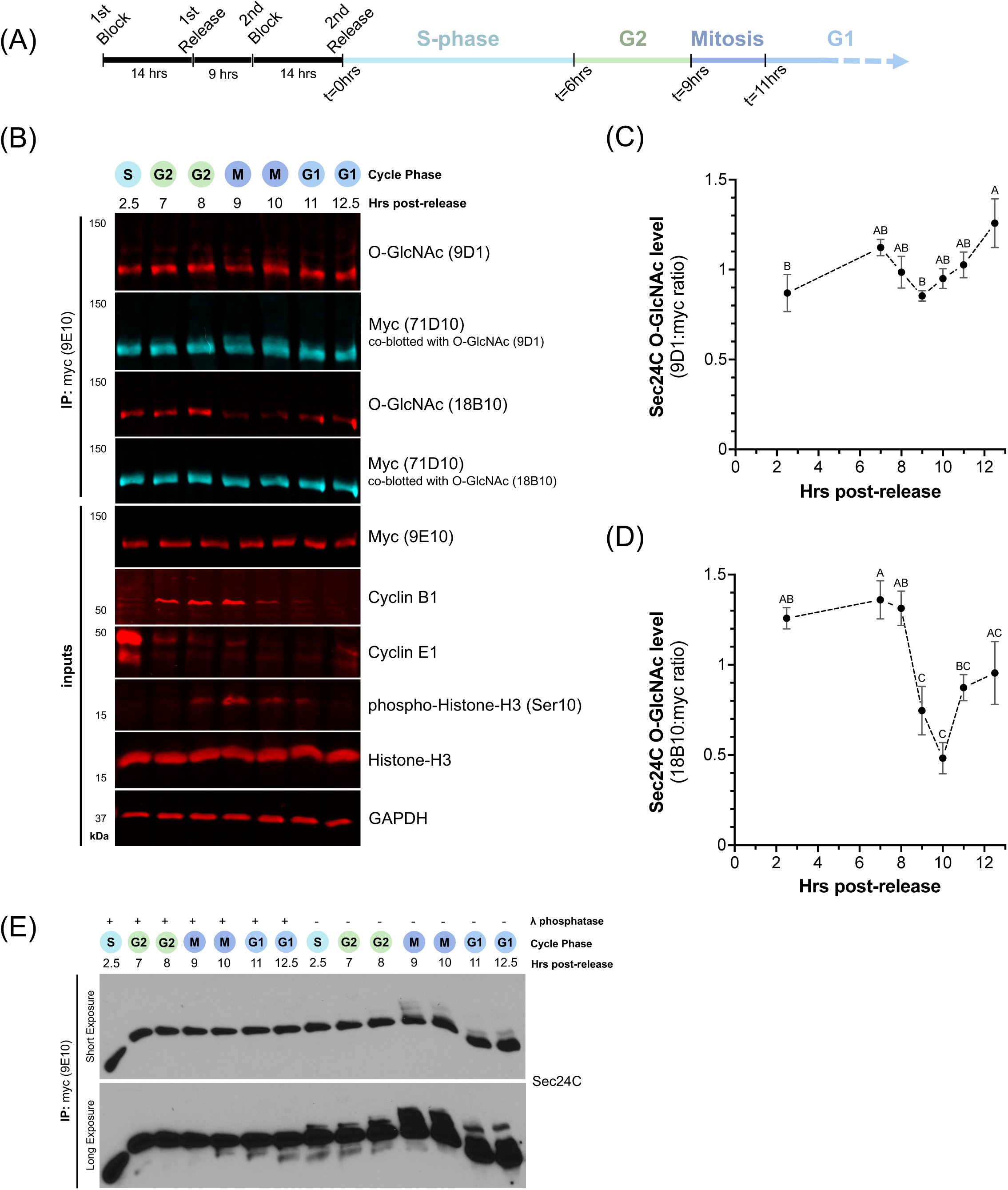
Sec24C is predominantly O-GlcNAcylated in interphase and phosphorylated in mitosis. (A) Treatment scheme. MH-Sec24C HeLa were synchronized using DTB to analyze PTMs at various cell cycle stages. Times listed after “2^nd^ release” denote approximate boundaries between cell cycle stages. (B) After DTB synchronization, MH-Sec24C cells were harvested at the indicated times, lysed, and analyzed by myc IP and IB. For O-GlcNAc quantification, membranes were stained simultaneously with anti-myc and anti-O-GlcNAc antibodies and secondary antibodies with distinct fluorophores, as indicated. (C, D) O-GlcNAc levels on Sec24C from synchronized cells at different cell cycle stages were quantified using two monoclonal anti-O-GlcNAc antibodies, 9D1 (C) and 18B10 (D). Data represent the mean of three or four biological replicates ± standard error of the mean (SEM), normalized to the mean of each replicate. Statistical significance was determined using one-way ANOVA with Tukey’s post-hoc test and is indicated using the compact letter display format (CLD). In CLD, any two data points with a letter in common in their labels (A, B, C, etc.) are statistically indistinguishable (*p* > 0.05), whereas any two data points without a letter in common are significantly different (*p* ≤ 0.05) from each other. (E) Cell lysates from (B) were subjected to myc IP, followed by treatment with λ phosphatase or mock conditions. Proteins were analyzed by Phos-Tag SDS-PAGE and IB to identify unique phosphoforms.

To examine Sec24C phosphorylation across the cell cycle, we employed Phos-Tag SDS-PAGE, which reports on a target protein’s phosphorylation **(62, 63)**. Manganese in the Phos-Tag gel retards the migration of phosphoproteins, allowing different phosphoforms to be detected as distinct bands of varying apparent molecular weight. This technique is particularly useful for Sec24C, as no anti-phospho-Sec24C antibodies are available, and pan anti-phosphothreonine and phosphoserine antibodies are non-specific (not shown). Phos-Tag analysis of cell cycle stage-enriched samples revealed that multiple putative Sec24C phosphoforms exist across phases (Fig. 1E). These distinct bands are authentic Sec24C phosphoforms because λ phosphatase treatment collapsed them to the predicted Sec24C molecular weight in untreated controls (Fig. 1E). At least one phosphoform was observed in S and G2 phases, two in G1, and three in mitosis (Fig. 1E). Notably, the proportion of phosphorylated Sec24C was greater in mitosis than in other phases (Fig. 1E). These results show that populations of phosphorylated Sec24C always exist in cycling HeLa cells but that mitotic Sec24C is more highly phosphorylated than interphase Sec24C and exists in at least one highly modified phosphoform not observed in other phases.

Together, these results demonstrate that Sec24C glycosylation and phosphorylation levels are responsive to cell cycle progression. While the extents of O-GlcNAcylation and phosphorylation are most divergent in mitosis, both modifications occur on Sec24C throughout the cell cycle, arguing against a simple reciprocal antagonism between them.

### Sec24C phosphorylation increases dramatically from mitotic entry through prometaphase

Mitosis is orchestrated by a complex and well-characterized cascade of phosphorylation events **(64–67)**. To determine whether Sec24C phosphorylation varies across the different sub-stages of mitosis and to confirm our DTB studies via another approach, we turned to nocodazole, a microtubule poison that traps cells in prometaphase via activation of the spindle assembly checkpoint **(68–71)**. Cells were treated for 0 to 20 hours with nocodazole or vehicle, and lysates were analyzed by IP-IB. Mitotic arrest was confirmed by IBs showing increasing cyclin B1 and pH3-Ser10 levels with continued incubation in nocodazole, compared to controls (Fig. 2A). Additionally, cells were assessed for rounding, which occurs during mitosis, by light microscopy. Nocodazole treatment for ≥16 hours resulted in nearly all cells in mitotic arrest (Fig. 2A).

**Figure 2.**
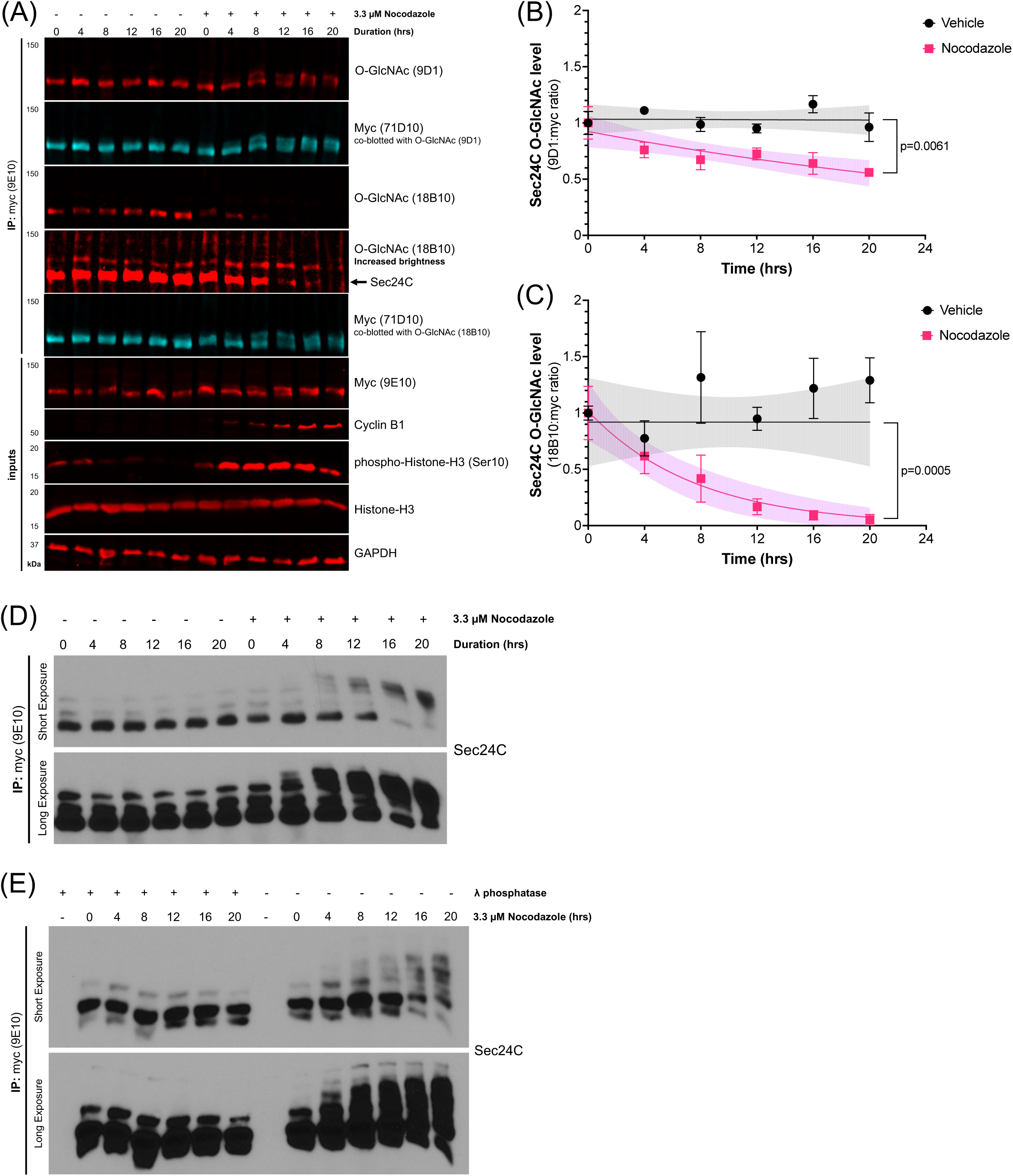
Mitotic arrest causes loss of Sec24C O-GlcNAcylation and increase in phosphorylation. (A) MH-Sec24C cells were treated with 3.3 µM nocodazole or DMSO (vehicle) as indicated and harvested, and lysates were analyzed by myc IP and IB. For O-GlcNAc quantification, membranes were stained simultaneously with anti-myc and anti-O-GlcNAc antibodies as indicated above. Black arrow indicates location of Sec24C. (B, C) O-GlcNAc levels on Sec24C from nocodazole-treated cells were quantified using two monoclonal anti-O-GlcNAc antibodies 9D1 (B) and 18B10 (C). Data represent the mean of three biological replicates ± SEM, normalized to the mean at t=0 for each replicate. Data were fit to one-phase decay models (lines) with 95% confidence intervals (shaded regions). Statistical difference between control and compound-treated samples was assessed by comparing decay models (k ± SEM, df) via unpaired t-test. (D) Cell lysates from (A) were subjected to myc IP and Phos-Tag SDS-PAGE, followed by IB. (E) Cell lysates from (A) were subjected to myc IP, λ phosphatase treatment (or mock treatment), and Phos-Tag SDS-PAGE.

Similar to our DTB results (Fig. 1B-D), Sec24C O-GlcNAcylation decreased in nocodazole-arrested mitotic versus control (asynchronous) cells (Fig. 2A). Moreover, this effect was more dramatic with prolonged arrest, with 18B10 signal on Sec24C nearly extinguished after 20 hours of treatment (Fig. 2A-B). We observed a similar trend with 9D1, albeit to a lesser extent (Fig. 2A,C), likely because different O-GlcNAc monoclonal antibodies are known to exhibit different amino acid sequence specificities **(72)**. Notably, nocodazole treatment led to the clear appearance of an upper Sec24C band (Fig. 2A), similar to mitotic samples from DTB-synchronized cells (Fig. 1B), and ≥16-hour treatment resulted in the near total loss of the lower band (Fig. 2A). The upper band is an authentic phosphoform of Sec24C, because it was eliminated by λ phosphatase (Fig. S3A). These results together indicate that prolonged prometaphase arrest causes the phosphorylation of nearly all Sec24C. While 9D1 blots revealed O-GlcNAcylation on both the upper, mitosis-associated and the lower, interphase-associated Sec24C bands, 18B10 only recognized the interphase-associated band, consistent with our observations that mitotic Sec24C is dramatically deglycosylated (Fig. 2A). Nocodazole had no impact on Sec24C protein levels (Fig. S3B).

Next, we used Phos-Tag analysis to examine mitosis-associated Sec24C phosphoforms. Consistent with the prior results, nocodazole treatment caused the appearance of higher phosphoforms in a time-dependent manner, with the number and abundance of phosphoforms in prometaphase-arrested cells exceeding those observed in DTB-synchronized mitotic cells (Fig. 2D-E). After 16-hour nocodazole treatment, when nearly all cells are prometaphase-arrested, the proportion of lower Sec24C phosphoforms and non-phosphorylated Sec24C decreased substantially (Fig. 2E). These results indicate that the entire cellular pool of Sec24C is rapidly and extensively phosphorylated upon mitotic entry and that the highly modified phosphoforms persist at least through prometaphase.

### Sec24C O-GlcNAcylation impacts its localization in prophase

O-GlcNAc and O-phosphate engage in well-documented cross-talk, sometimes competing for identical or nearby residues on substrates **(73–80)**. Our results showed that Sec24C is phosphorylated and deglycosylated upon mitotic entry (Fig. 1-2), raising the questions of whether Sec24C deglycosylation is required for its phosphorylation during mitosis and whether either PTM impacts Sec24C localization during cell division. To address these questions, we decoupled deglycosylation from mitotic entry by treating G2-synchronized cells with Thiamet-G (TG), a potent inhibitor of OGA, and examining Sec24C O-GlcNAcylation, phosphorylation, and localization in mitosis (Fig. 3A). To generate adequate numbers of mitotic cells for analysis, we included nocodazole treatment controls (Fig. 3A). Synchronization and mitotic arrest were confirmed as before (Fig. 3B). IP-IB analysis of Sec24C O-GlcNAcylation indicated that, compared to control, TG treatment in G2 increased Sec24C O-GlcNAcylation (Fig. 3B-D). However, nocodazole arrest in mitosis had no impact on Sec24C glycosylation in this context (Fig. 3C-D). Next, we tested whether enforced O-GlcNAcylation reduced mitotic Sec24C phosphorylation. IB demonstrated that the mitosis-associated Sec24C upper band was not diminished by treatment with TG, indicating that Sec24C phosphorylation in mitosis does not require deglycosylation (Fig. 3B). This conclusion was corroborated by Phos-Tag analysis, which showed no TG-dependent difference in Sec24C mitotic phosphoform number, pattern, or abundance (Fig. 3E). These results demonstrate that increased Sec24C phosphorylation in mitosis coincides with, but does not require, removal of O-GlcNAc, suggesting that the two PTMs modify distinct residues in this context.

**Figure 3.**
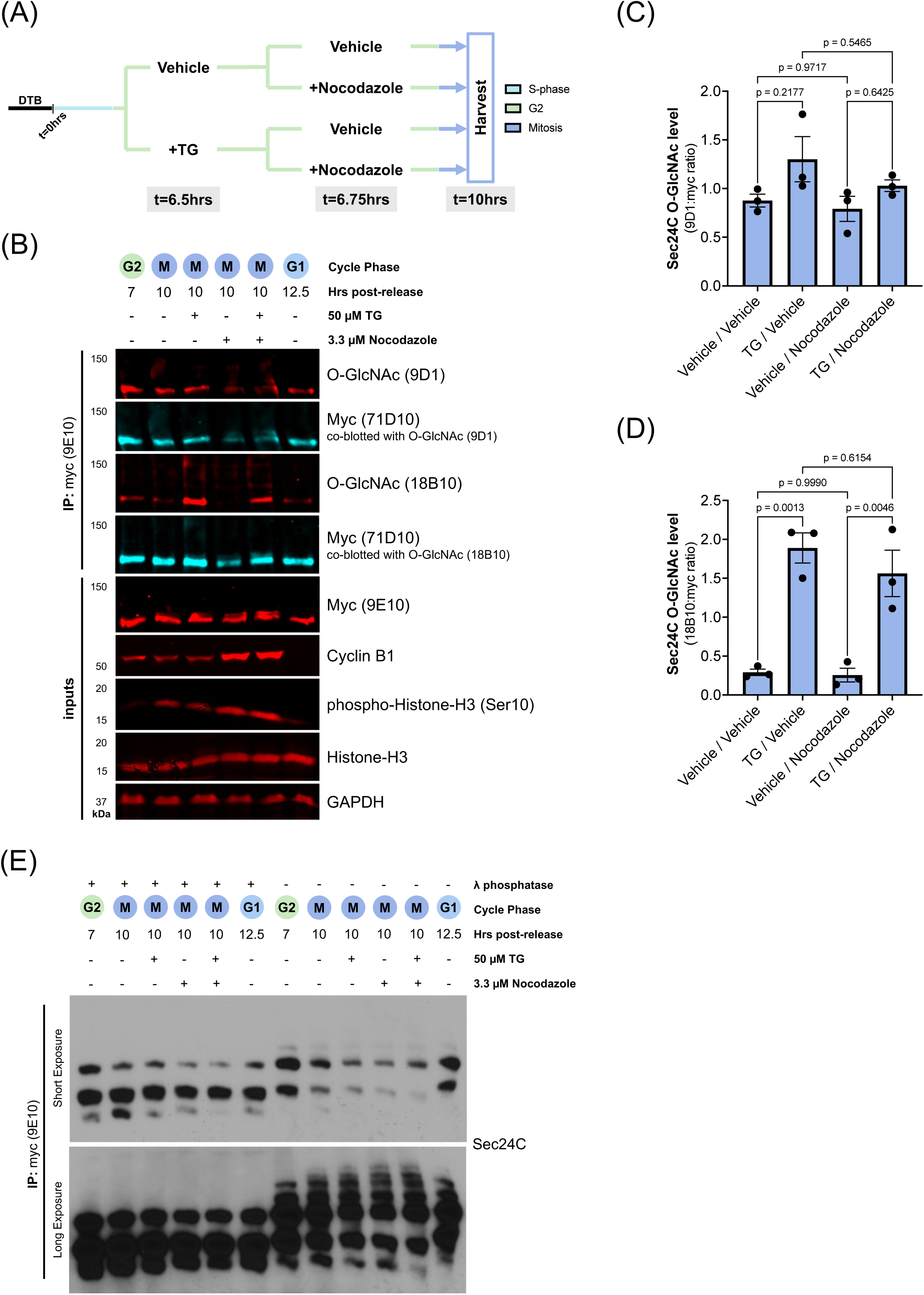
Mitotic Sec24C phosphorylation is independent of deglycosylation. (A) Treatment scheme. Cells were treated with either DMSO or 50 µM TG and either vehicle or 3.3 µM nocodazole at the indicated times post-release from DTB. (B) MH-Sec24C cells were treated as in (A), harvested, lysed, and subjected to myc IP and IB. t=7 and 12.5 hour samples serve as controls for cyclin and histone blots. For O-GlcNAc quantification, membranes were stained simultaneously with anti-myc and anti-O-GlcNAc antibodies as indicated above. (C, D) O-GlcNAc levels on Sec24C were quantified using two monoclonal anti-O-GlcNAc antibodies, 9D1 (C) and 18B10 (D), and compared across conditions. Data represent three biological replicates normalized to the mean of each replicate. Bars indicate means ± SEM. Statistical significance (*p* ≤ 0.05) was determined using one-way ANOVA with Tukey’s post-hoc test, with *p*-values displayed for comparisons of interest. (E) Cell lysates from (B) were subjected to myc IP, λ phosphatase treatment (or mock treatment), and Phos-Tag SDS-PAGE, followed by IB.

We next assessed whether Sec24C deglycosylation is required for its cytoplasmic dispersal during mitosis (Fig. 4). TG efficacy was reconfirmed by staining with RL2 (Fig. 4A-B), an anti-O-GlcNAc antibody preferred for its specificity in IF **(81)**, and successful synchronization and mitotic-arrest were confirmed by pH3-Ser10 staining (Fig. 4B). We then classified cells by cell cycle phase (Fig. S4A-B). Prophase, metaphase, anaphase, telophase, and mitotic exit (i.e., late telophase to early G1) could be distinguished via nuclear morphology and pH3-Ser10 intensity (Fig. S4A), but G1, S-phase, and early G2 could not, due to lack of changes in nuclear features and pH3-Ser10 levels throughout interphase (Fig. S4A-B). We note that there were insufficient anaphase cells observed to include in this analysis. Since Sec24C shifts from discrete puncta to a diffuse cytosolic distribution during mitosis, we quantified the percentage of total cell area occupied by Sec24C at different cell cycle stages and compared these values between TG-treated cells and control (Fig. 4D). (Nocodazole-treated samples were excluded from this analysis because nocodazole affects COPII subunit localization independent of its ability to induce mitotic arrest **(55, 82)**.) As expected, percent-area occupied by Sec24C greatly increased in mitotic stages, compared to interphase, with this increase beginning in the transition from late G2 to prophase and reaching its maximal point in metaphase (Fig. 4D). Interestingly, cells treated with TG did not undergo the same shift in Sec24C dispersal between late G2 and prophase (Fig. 4D-E), though dispersal at metaphase was indistinguishable between TG-treated cells and control (Fig. 4D). Consistent with these observations, the distribution of total Sec24C fluorescence across cell area reflected an identical trend (Fig. S4C). Taken together, these data show that removal of Sec24C O-GlcNAcylation prior to mitosis is required for its timely dispersal during mitosis but dispensable for its phosphorylation, and Sec24C phosphorylation upon mitotic entry may mediate its relocalization.

**Figure 4.**
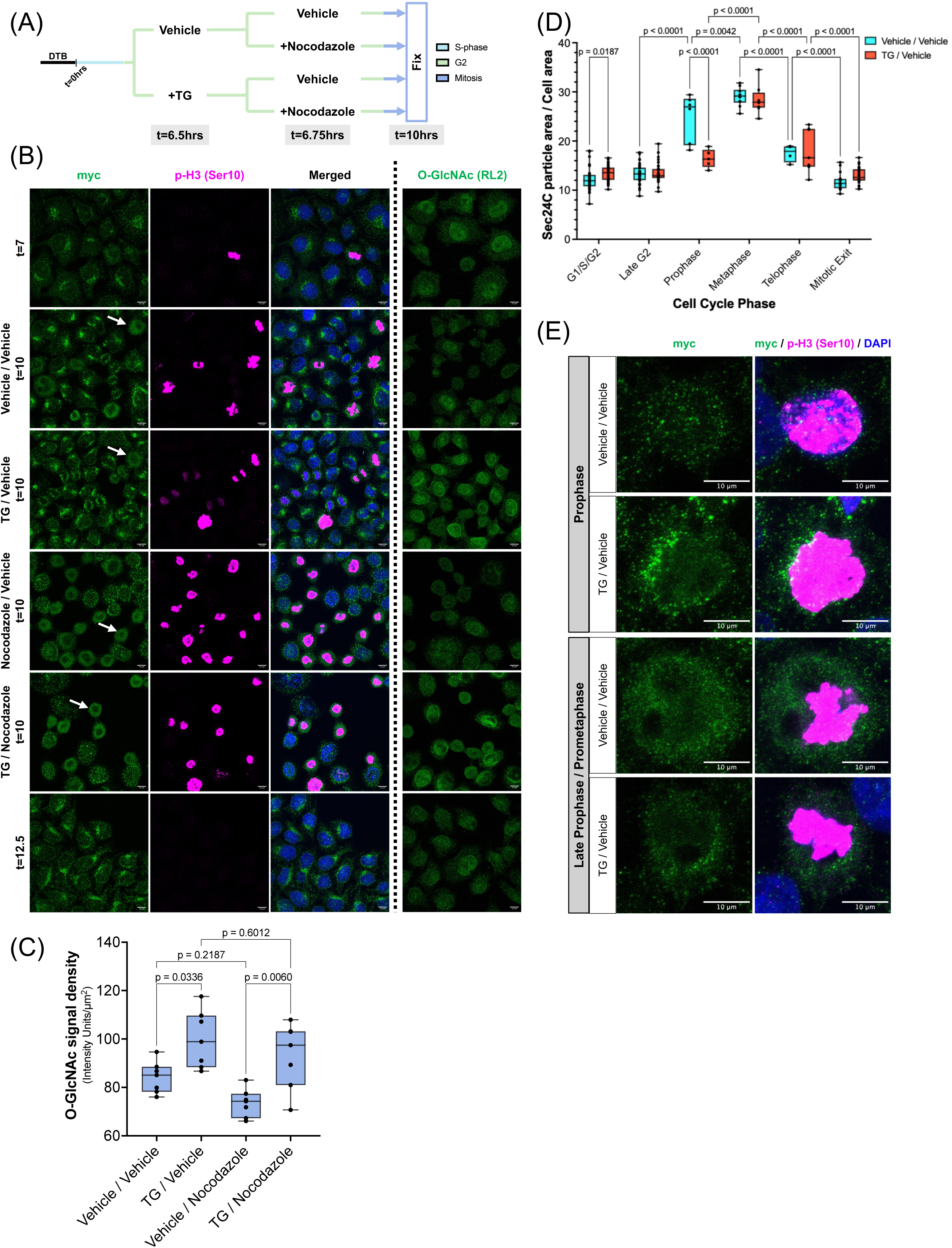
Sec24C O-GlcNAcylation affects dispersal in prophase. (A) Treatment scheme. Cells were treated with either DMSO or 50 µM TG and either DMSO or 3.3 µM nocodazole at the indicated times post-release from DTB. (B) After DTB synchronization and treatment as in (A), MH-Sec24C cells were analyzed by IF. White arrows indicate metaphase and/or nocodazole-arrested cells. O-GlcNAc (RL2) IF was performed on separate cover slips. Scale bar = 10 µm. (C) Quantification of total RL2 signal within a cell, normalized to cell area. Statistical significance (*p* ≤ 0.05) was determined using one-way ANOVA with Tukey’s post-hoc test, with *p*-values displayed for comparisons of interest. (D) For comparison of Sec24C localization across cell cycle phases and treatments, cells from the Vehicle/Vehicle and Vehicle/TG groups were categorized into phases based on nuclear morphology and pH3(Ser10) staining, and the percent area occupied by Sec24C particles per cell was compared across phase groups. Data are presented with medians indicated by solid lines, 25th-75th percentile range by boxes, and full range from minimum to maximum by whiskers. Statistical significance was assessed using two-way ANOVA followed by Tukey’s post-hoc test. Only comparisons between TG and Vehicle and consecutive cell cycle phases are shown; *p*-values > 0.05 are not displayed. (E) Representative images comparing localization of Sec24C in early and late prophase cells treated as in (A).

### Increased global phosphorylation promotes Sec24C dispersal from ERES

To test whether increased phosphorylation mediates the dispersal of Sec24C during mitotic entry, we treated cells with calyculin A, a potent inhibitor of protein phosphatase 1 (PP1) and protein phosphatase 2A (PP2A) **(83)**. Short treatment with calyculin A resulted in the appearance of multiple highly phosphorylated forms of Sec24C, as observed via Phos-Tag SDS-PAGE (Fig. 5A). Correlated with this, we noted a marked reduction in the number and brightness of juxtanuclear Sec24C puncta by IF (Fig. 5B). We observed a similar behavior by Sec31A (Fig. 5B). This result indicates that, upon treatment with calyculin, both Sec24C and Sec31A were more evenly distributed throughout the cytoplasm, as opposed to being concentrated at ERES, as in control cells (Fig. 5B). Indeed, calyculin A caused a significant decrease in intensity variance of Sec24C signal in interphase cells, indicating a more even distribution throughout each cell upon inhibitor treatment (Fig. 5C). This decrease in variance was similar to that observed when comparing untreated interphase and mitotic cells (Fig. 5C). Notably, calyculin treatment in interphase cells reduced variance to levels statistically indistinguishable from either treated or untreated mitotic cells, indicating that the dispersal of Sec24C initiated by mitosis could not be distinguished from the effect of calyculin by this measure (Fig. 5C). Additionally, we compared the dispersal effect of calyculin to that of mitosis on Sec24C by measuring the percent of total cell area occupied by Sec24C puncta, as before (Fig. 5D). Calyculin caused a significant increase in Sec24C dispersal by this measure as well, though this was increased still further upon mitotic entry (Fig. 5D). Taken together, these data are consistent with the hypothesis that increased Sec24C phosphorylation upon mitotic entry mediates its relocalization and that a mitosis-specific reduction of phosphatase activity and/or increase in kinase activity may play a role in this process.

**Figure 5.**
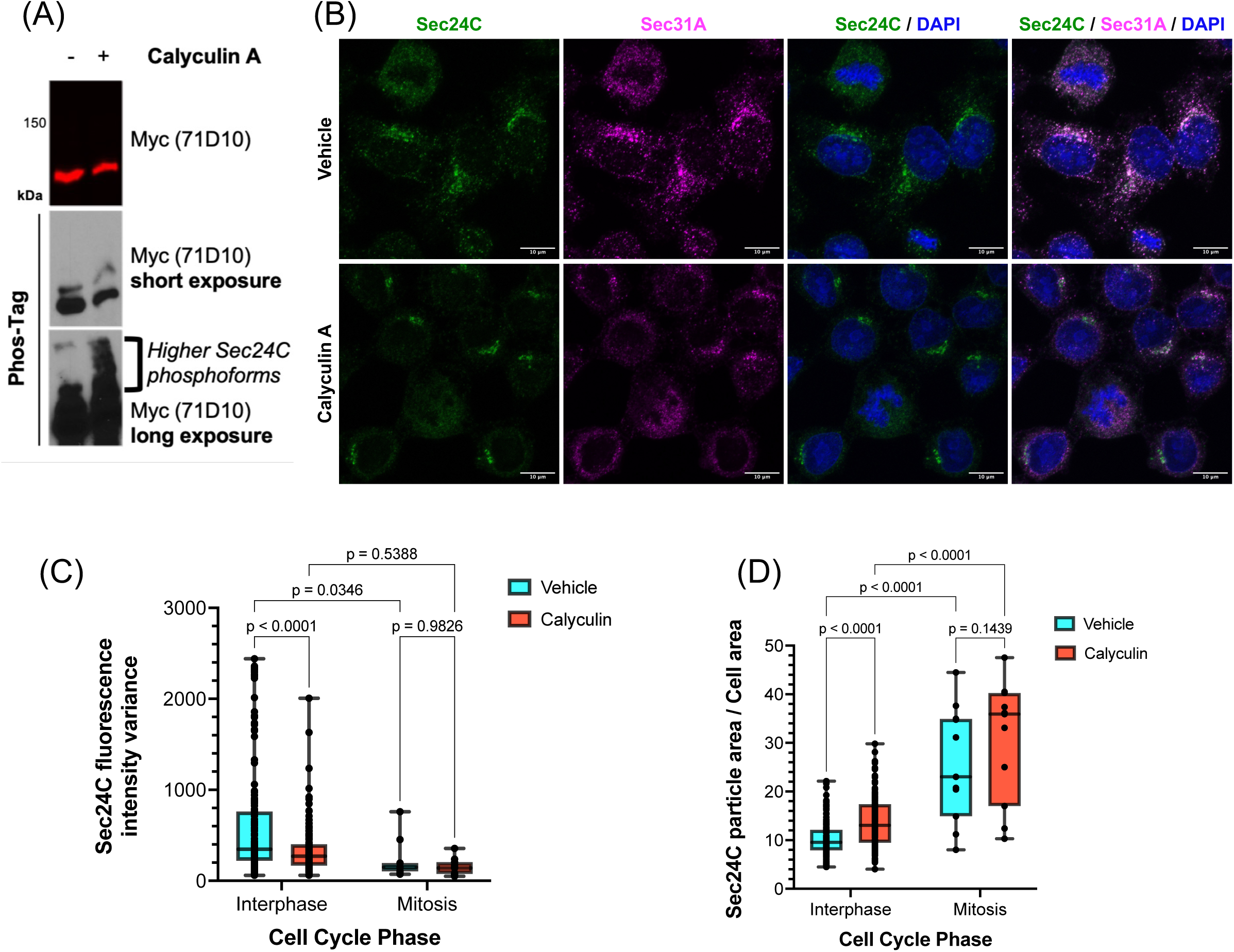
Increased Sec24C phosphorylation correlates with increased dispersal. (A) MH-Sec24C cells were treated with 10 nM calyculin A or DMSO for 30 minutes, harvested, lysed, and analyzed by standard or Phos-Tag IB. Black bracket indicates higher phosphoforms of Sec24C present in calyculin-treated sample. (B) Cells were treated as in (A) and analyzed by IF. Scale bar = 10 µm. (C, D) Sec24C signal from cells imaged in (B) was quantified to analyze localization. Cells were classified as interphase or mitotic based on nuclear morphology. (C) Variance of Sec24C fluorescence intensity per cell. (D) Area occupied by Sec24C particles within each cell as a percentage of total cell area. Data are presented with medians indicated by solid lines, 25th-75th percentile range by boxes, and full range from minimum to maximum by whiskers. Statistical significance was assessed using two-way ANOVA followed by Tukey’s post-hoc test. *p*-values are displayed for comparisons of interest.

### LC-MS/MS analysis of endogenous Sec24C from synchronized cells reveals novel glyco- and phosphosites

Our data suggest that site-specific O-GlcNAcylation and phosphorylation may regulate Sec24C localization and function across the cell cycle. To test this hypothesis systematically, we mapped PTMs on endogenous Sec24C in a cell cycle phase-specific manner. Cells were synchronized by DTB and harvested in G1, S, G2, or M, endogenous Sec24C was purified by tandem myc IP and Ni-NTA, and PTMs were analyzed by liquid chromatography and tandem mass spectrometry (LC-MS/MS) (Fig. 6A). We unambiguously identified one novel O-GlcNAc site at S205 and a novel phosphorylation site at T214 (Fig. 6B). Additionally, we observed evidence of phosphorylation at either T305 or S308 and simultaneous phosphorylation at three of the following four sites: T936, S937, T941, and T943 (Fig. 6B). In addition, upon reexamining Sec24C LC-MS/MS datasets that we published previously **(34)**, we observed phosphorylation at S888, which is a putative Sec24C phosphosite reported in a high-throughput screen for mitotic kinase targets **(84)** and a predicted site for the mitotic master regulator kinase, CDK1 **(85)** (Fig. 6B). In our system, we never observed evidence that any given Sec24C residue is both a phosphorylation and O-GlcNAcylation site, though prior reports indicate that S773 and T776, for example, can be modified by either PTM **(34, 84)**.

**Figure 6.**
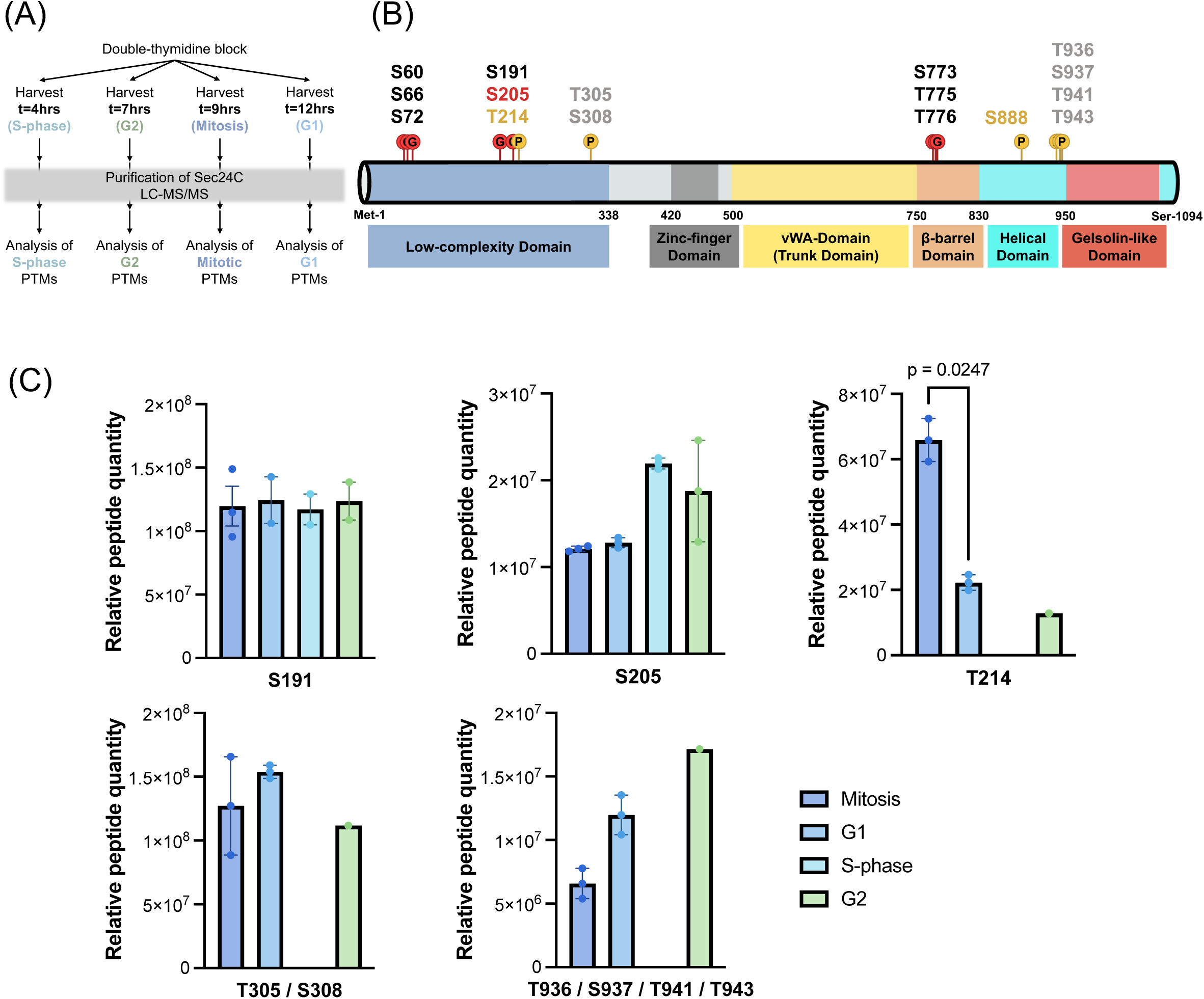
Identification of cell cycle phase-enriched PTMs of Sec24C. (A) Workflow for detecting cell cycle phase-enriched PTMs on endogenous human Sec24C. Harvest times are relative to time post-release from DTB. (B) Summarized MS results. Phosphosites are indicated by yellow circles with “P” and glycosites by red circles with “G.” Previously identified glycosites **(34)** are listed in black (S60, S66, S72, S773, T775, T776), novel glycosites in red (S205), and novel phosphosites in yellow or gray (T214, T305/S308, S888, T936/S937/T941/T943). Sites listed in gray are candidates because the specific phosphorylated residue could not be unambiguously assigned using the present MS data. (C) Quantification of peptides with specific PTMs using peak area integration from MS data. Peptide quantities were normalized to protein levels. Bar graphs show means ± SEM; individual data points represent biological replicates. Statistical significance was assessed using one-way ANOVA followed by Tukey’s post-hoc test, with significant comparisons (*p* < 0.05) indicated. Conditions where n ≤ 1 were excluded from statistical analyses.

None of the site-specific PTMs we observed was unique to a single cell cycle phase (Fig. 6C). However, quantifying and comparing the prevalence of each modified peptide among cell cycle phases revealed a significantly higher level of phopsho-T214 in mitosis than in G1 (Fig. 6C). This highlighted T214 as a particularly interesting PTM site for further exploration.

COPII trafficking and cell cycle progression are both highly regulated and highly conserved processes among all eukaryotes **(5, 86, 87)**. Therefore, functionally important, cell cycle phase-enriched PTMs on Sec24 proteins may also be broadly conserved. As a first step towards testing this hypothesis, we examined Sec24C residue conservation across a wide array of eukaryotic orthologs (Fig. S5A-B) and among the four human Sec24 paralogs (Fig. S5C). Generally, residues in the structured domains of Sec24C were more highly conserved than those in the low-complexity domain (LCD), though the conserved LCD O-GlcNAc site S191 is an exception (Fig. S5B-C). (We consider Ser↔Thr changes to be conserved for the purpose of analyzing O-linked PTMs.) Also of note, the β-barrel domain O-GlcNAc sites, S773, S776, and T776, were exceptionally well-conserved, as was the helical domain phosphorylation site, S888 (Fig. S5B-C). The high degree of evolutionary conservation of these residues is consistent with the hypothesis that their PTMs may be biologically significant and motivated us to assess their roles in the mitosis-associated regulation of Sec24C.

### Site-specific phosphorylation modulates the stability and localization of Sec24C

We tested the functions of selected Sec24C glyco- and phosphosites using a panel of high-priority mutants. First, we expressed unmodifiable Ala point-mutants of Sec24C in a Sec24C^−/-^ background **(34)** and compared their phenotypes to wild type (WT). Constructs expressing phosphomimetic (Ala to Asp) point-mutants were also used to assess the effects of site-specific phosphorylation. A compound Ala mutant, Sec24C^S773A-T775A-T776A^, abbreviated Sec24C^β-AAA^, was used to test the role of O-GlcNAcylation at PTM sites in the β-barrel domain **(34)**. Finally, we used compound Ala and Asp mutants to test the role of phosphorylation at the three simultaneously modified residues identified in the helical domain. Since these three phosphosites could not be unambiguously assigned among four possible residues, we selected and mutated T936, T941, and T943 to Ala or Asp (denoted Sec24C^He-AAA^ and Sec24C^He-DDD^, respectively) for initial studies.

Next, we analyzed mutants for differences in Sec24C protein levels, O-GlcNAcylation, and phosphorylation (Fig. 7, Fig. S6A, Fig. S6D-E) and for their ability to associate with the obligate binding partners Sec23A and Sec23B to ensure that Sec24C mutants were not grossly misfolded (Fig. S6B-C). Co-IP efficiency between Sec23 and each Sec24C mutant was unchanged, relative to WT (Fig. S6B-C). O-GlcNAc levels among mutants and WT were also very similar, as assessed by 9D1 IB (Fig. 7B). However, 18B10 IB revealed an increase in O-GlcNAcylation on Sec24C^S205A^ and Sec24C^S888D^ (Fig. 7C). Interestingly, mutation of S888 to alanine did not induce the same change (Fig. 7C). Additionally, Sec24C^β-AAA^ showed a drastic decrease in O-GlcNAcylation detected by 18B10 IB compared to WT, an observation that was statistically significant when comparing those two groups alone (Fig. S6A). No obvious differences were observed in the total phosphorylation of Sec24C mutants, compared to WT, except for Sec24C^S191A^, where a unique, highly phosphorylated upper band was sometimes observed (Fig. S6D-E).

**Figure 7.**
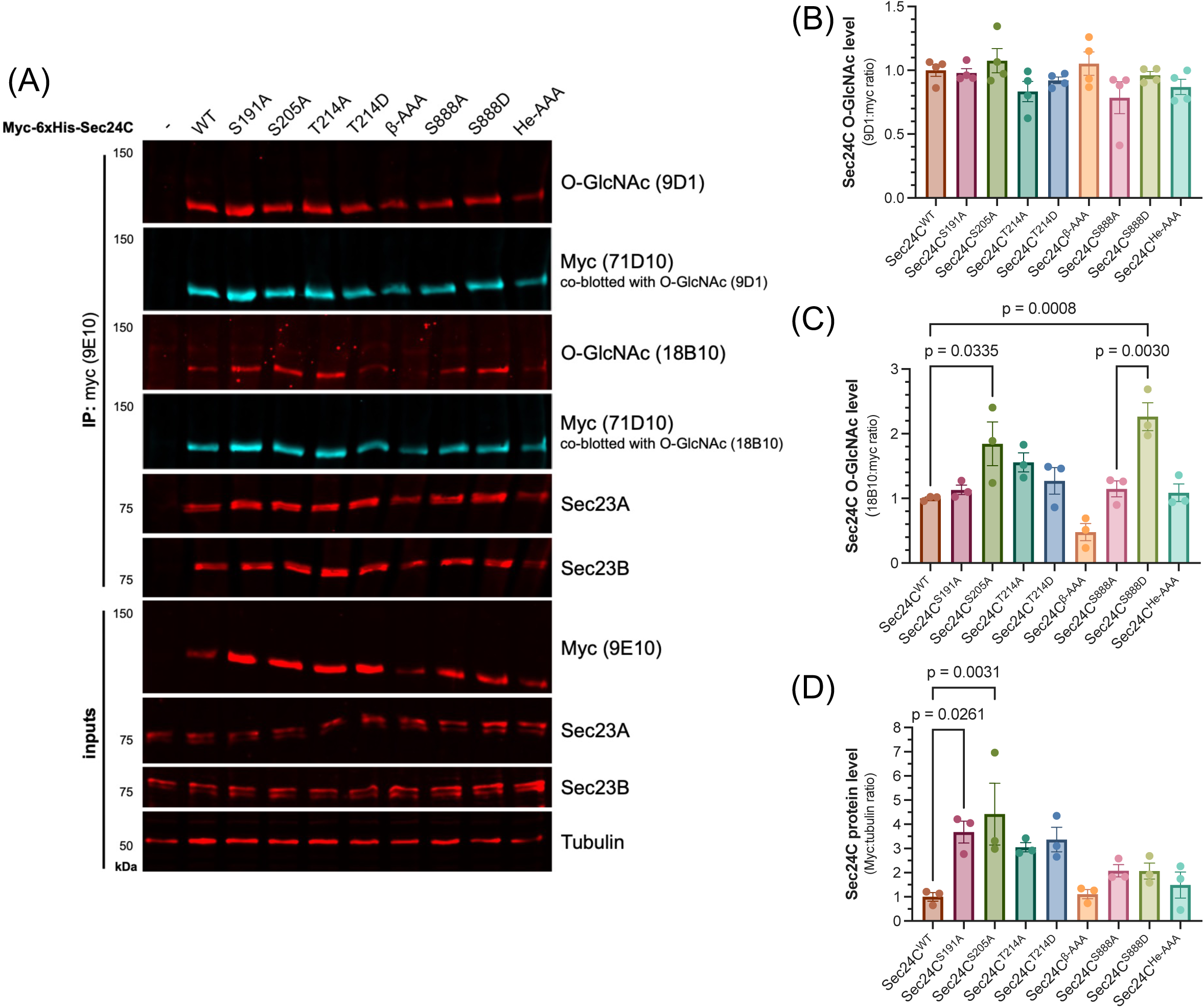
Point mutations reveal effects on Sec24C O-GlcNAcylation and protein level. (A) Sec24C-deficient HeLa cells (Sec24C KO) were transfected with empty vector (-), WT, single point mutants (S191A, S205A, T214A/D or S888A/D), or compound mutants (β-AAA or He-AAA) of Sec24C. 48 hours post-transfection, cells were harvested and lysed, and lysates were analyzed by myc IP and IB. For O-GlcNAc quantification, membranes were stained simultaneously with anti-myc and anti-O-GlcNAc antibodies as indicated above. (B, C) O-GlcNAc levels on Sec24C were quantified using two monoclonal anti-O-GlcNAc antibodies, 9D1 (B) and 18B10 (C). (D) Total Sec24C was quantified and normalized to tubulin for each sample. For (B-D), data points represent independent biological replicates (n=3 or 4) normalized to the WT sample mean for each replicate. Bars represent means ± SEM. Statistical significance was assessed by one-way ANOVA, and post-hoc comparisons were conducted between selected groups (WT versus all other constructs and Ala versus Asp mutants of the same residue) using Šídák’s method. Only statistically significant comparisons are shown.

Sec24C expression also varied across mutants (Fig. 7D). For example, the Sec24C^S191A^ and Sec24C^S205A^ glycosite mutants showed increased expression (Fig. 7D). Conversely, the expression of Sec24^He-DDD^ was extremely low, roughly 1/15^th^ the level of WT (Fig. 8A-B). Notably, however, expression of Sec24C^He-AAA^ was not different from WT (Fig. 8B). To determine whether Sec24C^He-DDD^ may be targeted for destruction, we performed a cycloheximide chase experiment to analyze its degradation rate in comparison to Sec24C^WT^ and Sec24C^He-AAA^ (Fig. 8C). Indeed, Sec24C^He-DDD^ was degraded much more rapidly than either WT or Sec24C^He-AAA^, while the degradation rate of Sec24C^He-AAA^ was statistically indistinguishable from that of WT (Fig. 8D). Together, these results suggest that compound phosphorylation at T936-T941-T943 may regulate Sec24C by marking it for degradation in response to unknown signals, a testable hypothesis for future work.

**Figure 8.**
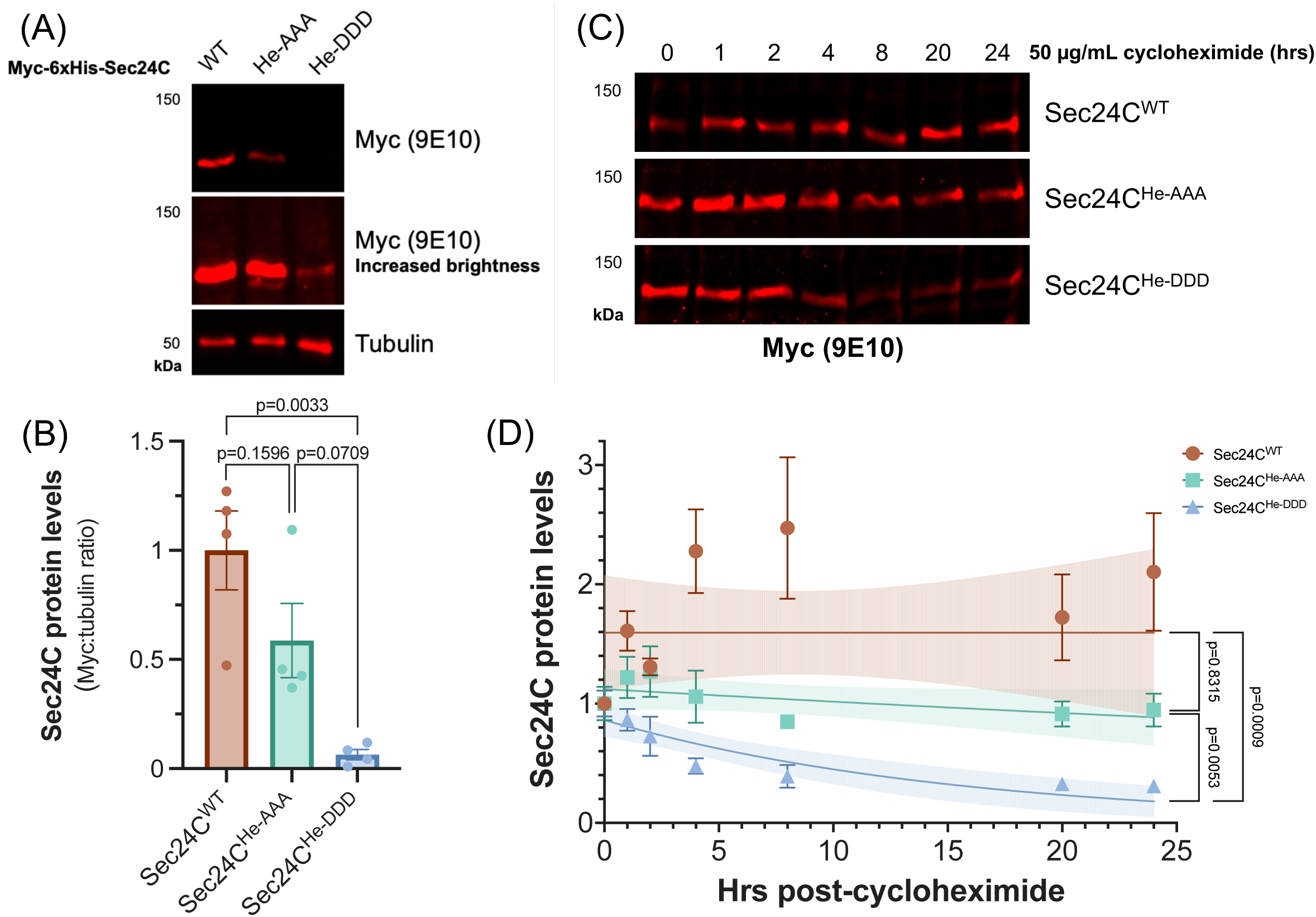
Helical domain phosphosites regulate Sec24C stability. (A) Sec24C KO cells were transfected for 48 hours with WT, He-AAA, or He-DDD Sec24C, and lysates were analyzed by IB. (B) Levels of Sec24C (normalized to tubulin) were quantified for each sample. Data points represent independent biological replicates (n=4) normalized to the WT mean for each replicate. Bars represent mean ± SEM. Statistical significance was assessed by one-way ANOVA with Tukey’s post-hoc test. Only statistically significant comparisons are shown. (C) Sec24C KO cells were transfected for 24 hours and then treated with 50 µg/mL cycloheximide as indicated. Lysates were analyzed by IB. (D) Quantification of change in Sec24C (myc) level. Data points represent the mean of three biological replicates ± SEM, normalized to the mean at t=0 for each replicate. Data were fit to one-phase decay models (lines) with 95% confidence intervals (shaded regions). Statistical difference between Sec24C constructs was assessed by comparing decay models (k ± SEM, df) via one-way ANOVA with Tukey’s post-hoc test.

Lastly, to ascertain whether any of the identified PTMs might mediate Sec24C mitotic dispersal, we examined mutant localization by IF. Imaging results corroborated the IB data, showing a clear reduction in the intensity of Sec24C^He-DDD^ versus WT and Sec24C^He-AAA^ puncta (Fig. 9A). Moreover, Sec24C^He-DDD^ differed in how puncta were distributed, with no clear juxtanuclear localization, as is observed in WT. Instead, Sec24C^He-DDD^ showed a more uniform cytosolic distribution, reminiscent of the mitotic pattern of endogenous WT Sec24C (Fig. 9A). To further confirm these observations, we assessed the ratio of Sec24C puncta overlapping with Sec16A-marked ERES for WT versus Sec24C^He-DDD^. As expected, Sec24C^He-DDD^ showed a significant reduction in ERES-overlap, compared to Sec24C^He-AAA^ or WT, indicating an increase in its cytosolic dispersal (Fig. 9B). We next measured the distance between Sec24C puncta and nearest ERES as a more precise readout for cytosolic dispersal. For WT and all other mutants tested, the median distance to the closest ERES was 0 µm, whereas for Sec24C^He-DDD^, it was ∼35 nm (Fig. 9C). Mean distance to ERES also significantly differed between Sec24C^He-DDD^ and WT but not between Sec24C^He-AAA^ and WT (Fig. 9D). IF analysis additionally revealed a small but significant reduction in the mean distance to ERES of Sec24C^T214A^ in comparison to both Sec24C^T214D^ and WT (Fig. 9D). We also measured the mean and median volumes of ERES-localized and cytosol-localized Sec24C puncta, respectively, as metrics of COPII subunit recruitment and observed minor differences between some mutants and WT (Fig. S7A-D). For example, the volumes of cytosolic Sec24C^He-DDD^ puncta were significantly greater than those of WT and Sec24C^He-AAA^ (Fig. S7A-B). All together, these data provide strong evidence for the biological importance of both O-GlcNAcylation and phosphorylation in regulating the intracellular dynamics of Sec24C, including its localization, recruitment, stability, and response to cell cycle phase-specific and/or other stimuli.

**Figure 9.**
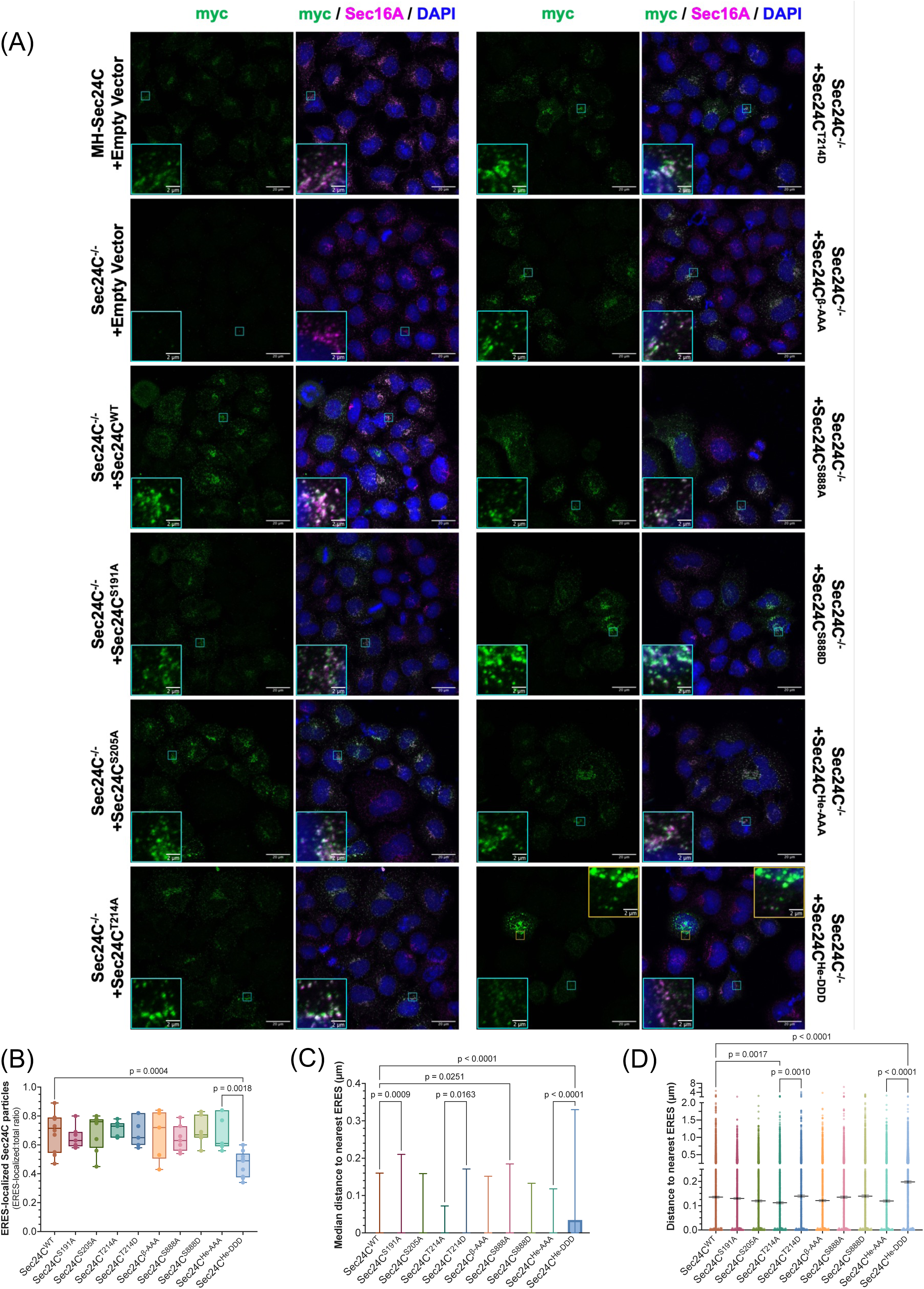
Sec24C PTMs regulate its localization and recruitment to ERES. (A) Sec24C KO HeLa cells were transfected as indicated for 24 hours and analyzed by IF. Scale bar = 10 µm; inset scale bar = 2 µm. (B) The ratio of Sec24C particles overlapping Sec16A-marked ERES was measured for each IF image. Data are presented with medians indicated by solid lines, 25th-75th percentile range by boxes, and full range from minimum to maximum by whiskers. Differences between means were assessed by one-way ANOVA and Šídák’s multiple comparisons test. (C) Median and (D) mean separation of Sec24C particles from nearest Sec16A-marked ERES. (C) Medians are shown as bars ± interquartile range. (D) Means are displayed as black lines ± SEM within scatter plots showing all values. For each replicate (n=3 to 5) and Sec24C construct tested, 998 Sec24C particles were measured. Differences between medians were assessed by Kruskal-Wallis test, followed by Dunn’s multiple comparison. Differences between means were assessed by one-way ANOVA and Šídák’s multiple comparisons test. Comparisons were only computed for select combinations (WT versus all other constructs and Ala versus Asp mutants of the same residue). Only statistically significant comparisons are shown.

## Discussion

Given the volume and diversity of proteins trafficked through the ER, understanding the regulation of secretory transport is critical for our understanding of basic eukaryotic cell biology and for insight into specific bone and bleeding disorders **(26–29, 88–91)** and broader topics ranging from neuronal health to viral infection **(23, 92, 93)**. Here, we characterized the cell cycle-dependent regulation of Sec24C, a core COPII subunit, by reciprocal O-GlcNAcylation and phosphorylation. We demonstrated that endogenous Sec24C is both phosphorylated and O-GlcNAcylated throughout the cell cycle but that O-GlcNAcylation peaks in G2 and reaches a minimum during mitosis, when phosphorylation reaches its maximum (Fig. 10). Our results differ somewhat from those of a prior study in which the authors could detect neither Sec24C glycosylation in mitotic cells nor Sec24C phosphorylation in interphase cells **(43)**. However, the authors used a different anti-O-GlcNAc antibody than we did and did not use Phos-Tag SDS-PAGE or MS analysis, which likely explains the apparent discrepancies. Indeed, Phos-Tag SDS-PAGE was a particularly useful tool, revealing not only cell cycle-dependent changes in Sec24C phosphorylation but also the fact that a minor fraction of total Sec24C is always phosphorylated. Future studies will focus on the function of this phosphoform, as its persistence throughout the cell cycle may suggest an important role in COPII trafficking or regulation.

**Figure 10.**
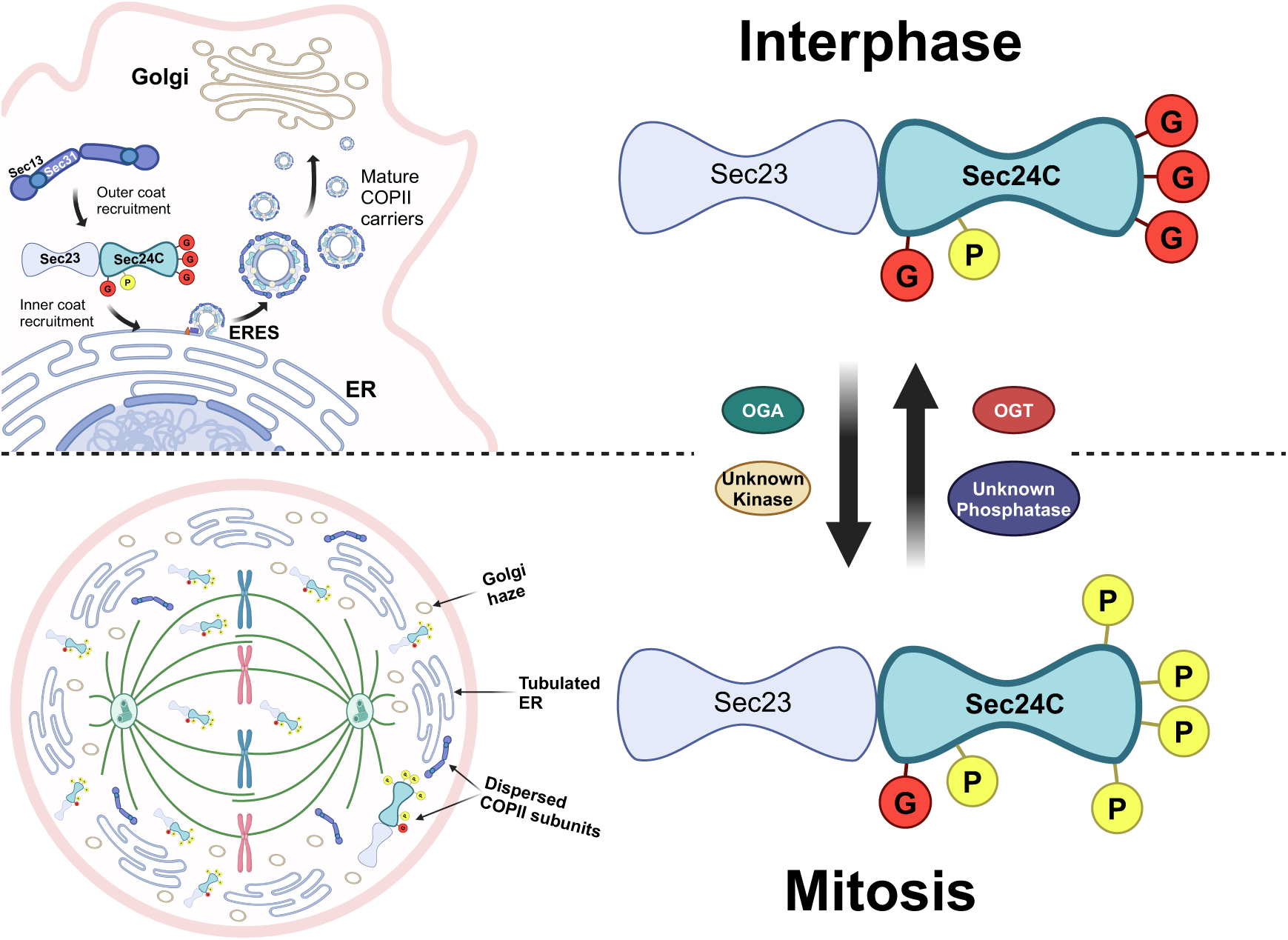
Working model of Sec24C PTMs and activity in interphase and mitosis. In interphase, Sec24C is more heavily O-GlcNAcylated, with lower-level phosphorylation. Interphase Sec24C interacts with Sec23, Sar1, and the outer coat proteins Sec13 and Sec31 at Sec16-marked ERES to form functional COPII carriers. In mitosis, COPII carrier components disperse from Sec16-marked ERES. This relocalization correlates with a rapid increase in Sec24C phosphorylation and decrease in Sec24C O-GlcNAcylation, likely at distinct residues.

On the other hand, our data do support a more complex form of crosstalk between O-GlcNAc and O-phosphate during cell cycle progression. In a uniformly mitotic population of cells, we observed a near-total elimination of Sec24C O-GlcNAcylation, as detected by the 18B10 antibody, which coincided with a dramatic increase in the number and intensity of observed phospho-forms (Fig. 2). We note that prometaphase arrest was achieved with nocodazole, a pleiotropic compound that is well-documented for its ability to disrupt the Golgi into mini-stacks **(55)**. We cannot rule out the possibility that Golgi disruption *per se* impacted COPII PTMs in our system, independent of cell cycle effects. However, our data show increases in Sec24C phosphorylation and higher phosphoforms that strongly correlate with increasing mitotic index (Fig. 2) and agree with results from DTB experiments (Fig. 1), where the Golgi is intact. Whether nocodazole or Golgi disruption affects COPII regulation through PTMs in a cell cycle-independent manner will require future studies.

Our studies posed two key questions: 1) Are Sec24C mitotic phosphorylation and deglycosylation causally linked or simply correlated? and 2) Do dynamic PTMs influence the mitotic dispersal of Sec24C? It has long been known that Sec24 and other COPII proteins change from their mostly perinuclear localization in interphase to adopt a more uniform, cytosolic distribution in mitosis **(44, 45)**. Furthermore, relocalization coincides with both a cessation in COPII trafficking and the dissolution of the Golgi, with some authors hypothesizing that the former at least partially mediates the latter **(94–96)**. The upstream driver of mitotic Golgi dissolution is debated, but evidence indicates that dispersal depends on mitotic kinases modifying the proteins that maintain the Golgi ribbon and stacks **(48–51, 53, 56, 57)**. These kinases include CDK1 and Plk1, the “master regulators” of mitosis, which have also been predicted to modify Sec24C, to thus-far unknown effect **(84, 85)**. Dudognon et al. first proposed that mitotic phosphorylation of Sec24C may cause its dispersal and contribute to the cessation of COPII trafficking during mitosis **(43)**. Indeed, the authors found that mitotic Sec24C could not interact with membranes in an *in vitro* recruitment assay **(43)**. Interestingly, a recent study found that an increase in phosphorylation of the COPII-related protein TANGO1 during mitosis was at least partially responsible for the mitotic dissolution of ERES in HeLa cells **(47)**. TANGO1 phosphorylation alone could not entirely explain the phenomenon, however, leaving open the possibility that concurrent phosphorylation of multiple COPII proteins promotes ERES dissolution in mitosis. Because the responsible kinase(s) has not been identified, we tested the potential role of Sec24C deglycosylation by blocking the activity of OGA prior to mitotic entry and quantifying the impact on Sec24C phosphorylation and dispersal (Fig. 3). We posited that if mitotic deglycosylation is required for phosphorylation, preventing deglycosylation would also prevent phosphorylation. Instead, we found that mitotic Sec24C phosphorylation still occurred despite high levels of O-GlcNAc (Fig. 3). This result implies that pre-mitotic O-GlcNAc and mitotic O-phosphates do not compete to modify the same residues on Sec24C, and the former does not preclude the latter.

On the other hand, OGA inhibition affected the cytosolic dispersal of Sec24C during prophase, causing a delay into metaphase but not a complete block (Fig. 4). The limitations of our experimental systems prevented us from directly demonstrating that dispersal is dependent on mitotic phosphorylation, but other data are consistent with this hypothesis. For example, increasing Sec24C phosphorylation by inhibiting PP1 and PP2A potentiated Sec24C dispersal (Fig. 5). If mitotic phosphorylation is the cause of Sec24C dispersal, an O-GlcNAc-dependent delay in dispersal may be due to retained glycosylation hindering the addition of mitotic phosphorylation. In fact, a similar phenomenon has been described for H3. O-GlcNAcylated H3 is deglycosylated near mitotic entry, and this deglycosylation promotes its phosphorylation **(97)**. H3 phosphorylation, in turn, is necessary for its mitotic activity **(97)**. While H3 deglycosylation is not required for its normal mitotic activity, blocking deglycosylation perturbs the timing of mitosis through its effect on H3 phosphorylation **(97)**. It will be important for future studies to determine whether and which specific Sec24C PTM sites are required for the effects we observed, particularly since chemical inhibitors of OGA or phosphatases impact many substrates simultaneously.

Our MS studies provide the first systematic analysis of PTMs on Sec24C expressed at WT levels from its endogenous locus. Though we did not observe any glycosylations exclusive to interphase or phosphorylations exclusive to mitosis, as predicted by prior literature **(43)**, we did observe other cell cycle phase-enriched effects (Fig. 6). For example, phospho-T214 peptides were about three times more abundant in mitosis-enriched than G1-enriched samples, revealing at least one phase-dependent phosphorylation site (Fig. 6). We speculate that more cell cycle phase-specific PTMs of Sec24C may remain to be discovered. Because PTMs are substoichiometric and often short-lived, many modifications are challenging to detect **(98–101)**. Furthermore, mitosis-specific phosphorylation on Sec24C is expected to occur in the short timeframe between prophase and telophase, and its glycosylation is also dynamic, adding to the difficulty of capturing and identifying all relevant sites **(98–101)**. We also observed incomplete peptide coverage of the Sec24C sequence across our combined LC-MS/MS replicates, which may have prevented identification of some PTMs. For example, as noted, phosphorylation on S888 was identified by reanalysis of previously published data, whereas our current MS data lacked coverage of S888. For these reasons, our data do not preclude the existence of additional phase-enriched or -specific PTM sites on Sec24C.

Mutagenesis studies based on our MS data also revealed new information about Sec24C PTMs. For instance, we observed an approximately 50% reduction in O-GlcNAc on Sec24C^β-AAA^ , compared to WT, indicating that these residues may be among those deglycosylated upon mitotic entry (Fig. 7, Fig. S6A). On the other hand, we observed that loss of O-GlcNAcylation at S205 caused increased overall levels of Sec24C O-GlcNAcylation (Fig. 7A, Fig. 7C). This phenomenon hints at the complex regulatory mechanisms underlying Sec24C modification – though S205 is not itself a major glycosite recognized by 18B10, our data indicates that glycosylation at this residue may modulate O-GlcNAcylation at other major Sec24C O-GlcNAc sites. A similar phenomenon has been observed for p65, where authors found that loss of glycosylation at T305 led to an increase in overall glycosylation through T352 **(102)**. Interestingly, an increase in overall Sec24C O-GlcNAcylation was also observed when the S888 phosphosite was mutated to Asp but not when mutated to Ala, suggesting that phosphorylation at this site increases O-GlcNAc at other site(s) (Fig. 7A, Fig. 7C) **(73, 79)**. To unravel the cause and function of this crosstalk, future studies will employ additional mutagenesis to identify the O-GlcNAc sites affected by S888 phosphorylation and biochemical studies to identify the responsible kinase (e.g., CDK1).

Lastly, we found that simultaneous mutation of helical domain residues T936, T941, and T943 to phosphomimetic Asp induced rapid degradation of Sec24C (Fig. 8). This mutant did not display typical perinuclear ERES localization, instead usually appearing evenly dispersed and occasionally as large and bright puncta (Fig. 9). Phosphorylation at these sites may constitute a regulatory mechanism for rapid degradation of Sec24C under unknown circumstances. Future studies should focus on elucidating the degradation mechanism, identifying upstream cues that induce these phosphorylations, and pinpointing the responsible kinase(s).

Taken together, our results demonstrate the dynamic modification of the core COPII inner coat subunit, Sec24C, by both phosphorylation and O-GlcNAcylation across the cell cycle. Beyond regulating trafficking, these PTMs in concert may contribute to Sec24C stability and redistribution during mitosis and contribute to the larger program of partitioning the endomembrane machinery between daughter cells after cell division. Our work also adds to a growing body of evidence that phosphorylation and O-GlcNAcylation of COPII subunits occur in response to cell signals and regulate aspects of the early secretory pathway, potentially with distinct effects on different Sec24 paralogs. Future work will dissect the functional importance of these PTMs and the crosstalk between them in the enigmatic behavior of COPII during mitosis.

## Experimental Procedures

### Materials

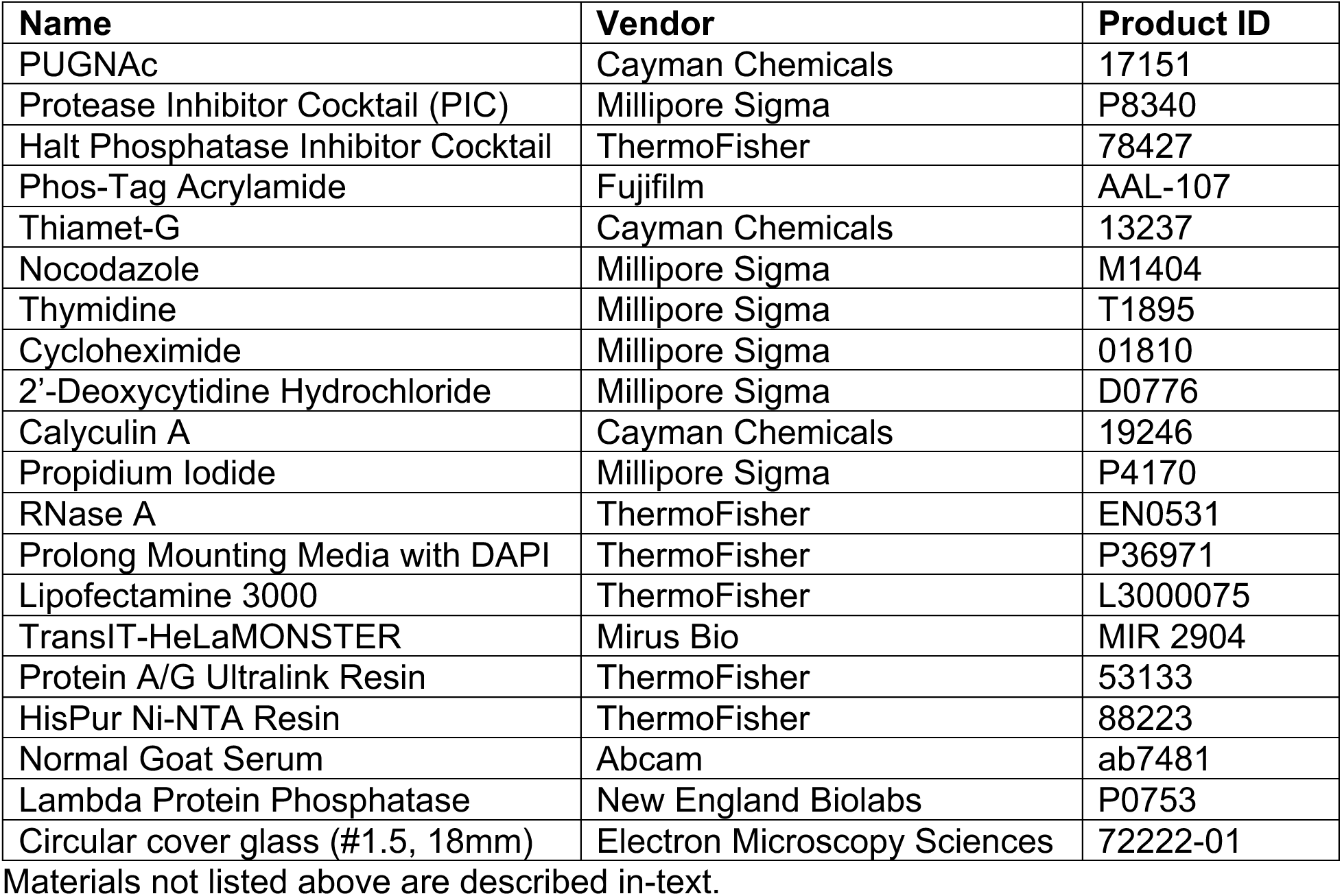

### Antibodies

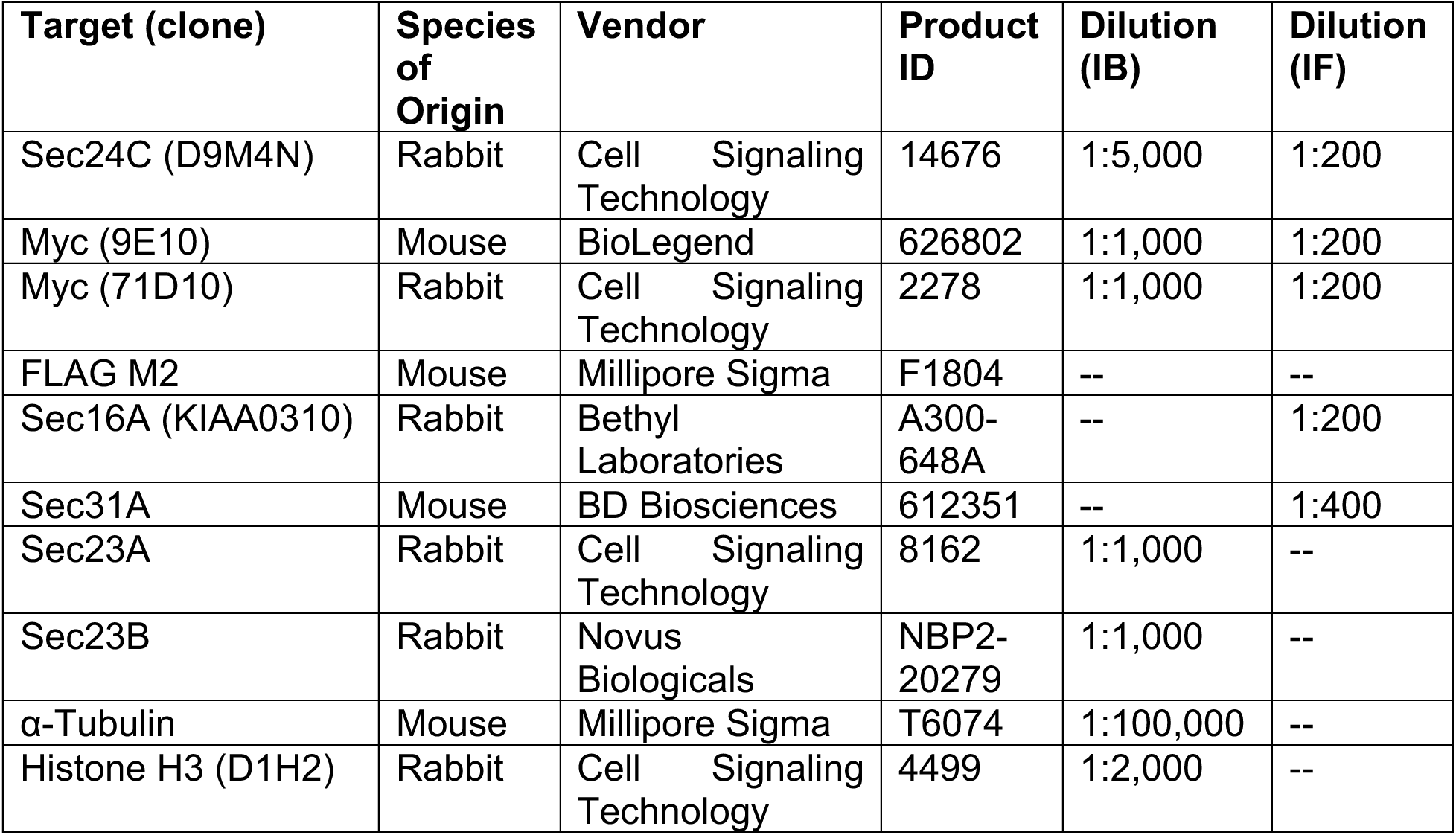
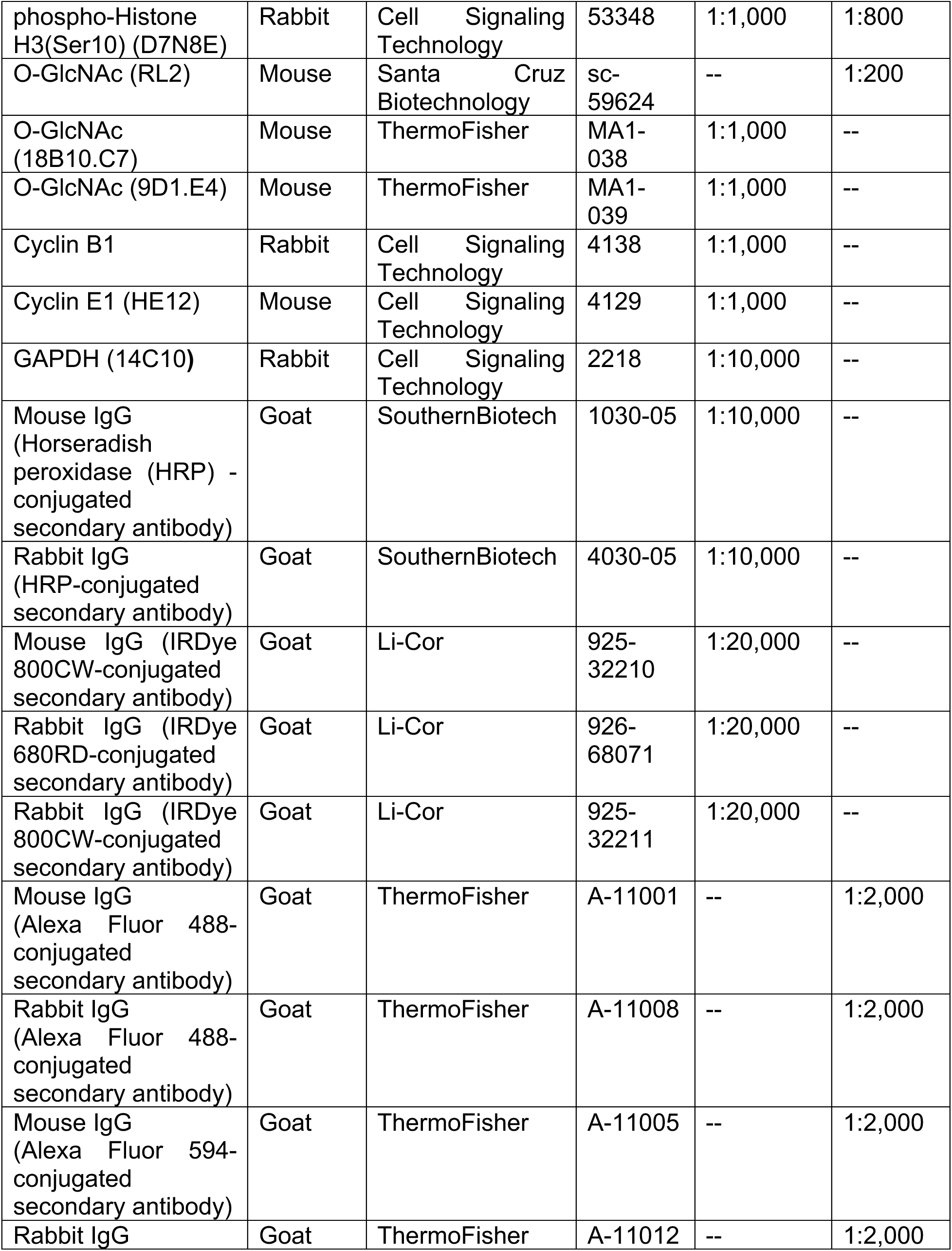
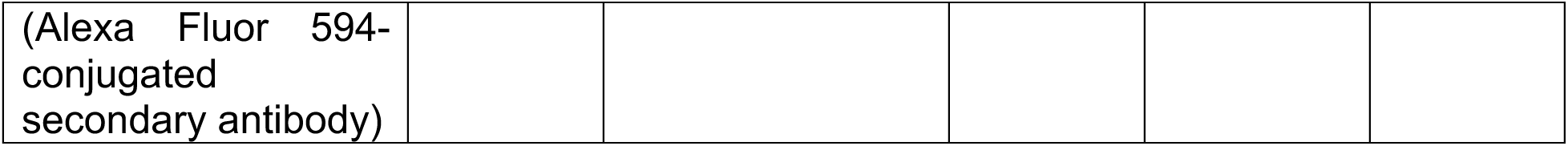

### DNA Constructs and CRISPR-Tagging of Endogenous Sec24C

Subcloning of N-terminally myc-6xHis-tagged WT Sec24C into pLenti CMV/TO Puro DEST (670–1) (Addgene, 17293) was previously described **(34)**. S191A and β-AAA mutants were previously prepared by mutagenesis of the WT construct **(34)**. S205A, T214A, and T214D were prepared from WT by site-directed mutagenesis. He-AAA and He-DDD mutants were prepared by subcloning small gene blocks encoding for the relevant mutations into the pLenti backbone via Gibson assembly. S888A and S888D mutants were prepared by Genewiz PriorityGene service by restriction cloning gene blocks containing the relevant mutations into the pLenti backbone using XmaI and AscI restriction enzymes. All in-lab mutagenesis was done via PCR using Phusion Hot Start II DNA polymerase (ThermoFisher, F549S), followed by DpnI (New England Biolabs [NEB], R0176S) digestion to eliminate template DNA, excision of appropriate bands after agarose gel electrophoresis, purification from the gel using a DNA recovery kit (Zymo Research, D4002), and ligation using NEBuilder HiFi DNA Assembly Master Mix (NEB, E2621). Where Gibson assembly was required, NEBuilder HiFi DNA Assembly Master Mix was used on purified PCR products at a 1:3 vector:insert mass ratio as determined by NEBioCalculator (https://nebiocalculator.neb.com/#!/ligation). Purified and assembled DNA products were transformed into NEB Stable Competent E. coli (NEB, C30401), after which, bacteria were streaked onto appropriate LB agar plates, and single colonies were picked for maxi-prep with ZymoPURE II Plasmid Purification Kit (Zymo Research, D4202) and Sanger sequencing. Mutagenesis primers were determined using NEBaseChanger (https://nebasechanger.neb.com/). Gibson assembly primers were designed using NEBuilder Assembly Tool (https://nebuilder.neb.com/#!/).

The endogenously myc-6xHis-tagged “MH-Sec24C” clonal cell line was produced using a CRISPR-Cas9 system and homologous recombination, as previously described **(59)** with some modifications. Briefly, a guide RNA (gRNA) targeting the N-terminus of Sec24C was designed using the Synthego Knockout Guide Design tool (https://design.synthego.com/<u>#/</u>), then subcloned into pSpCas9(BB)-2A-GFP(Px458) (Addgene, 48138) by the Duke Functional Genomics Core Facility. To incorporate the myc-6xHis tag into Sec24C at its endogenous locus, an ∼1000 bp homology-directed repair (HDR) template was designed such that its 5’ half was homologous to the sequence 5’ of the spCas9 cut site, and its 3’ half homologous to the sequence 3’ of the spCas9 cut site. The PAM site in the HDR template was mutated to avoid recognition and cutting by spCas9. The HDR template was purchased as a gene block from IDT and subcloned into AAVS1_Puro_PGK1_3xFLAG_Twin_Strep (Addgene, 68375) via Gibson assembly as described above. Parental HeLa cells were co-transfected using TransIT-HeLaMONSTER with either spCas9-GFP/gRNA and an empty repair template (negative control), or spCas9-GFP/gRNA and the HDR template at a 1:2 mass ratio. 24 hours post-transfection, cells were prepared for sorting following “Single Cell Sorting” and “Sample Preparation Guidelines for Cell Sorting” protocols from the UWCCC Flow Cytometry Laboratory (https://cancer.wisc.edu/research/resources/flow/). Using a Sony SH800 cell sorter (Duke Cancer Institute, Flow Cytometry Core), cells were selected for high GFP expression and seeded at a concentration of one cell per well of a 96-well plate. Once expanded sufficiently, surviving single cell-derived clones were tested by IP/IB for expression of myc-6xHis-tagged Sec24C, co-IP with Sec23A, and colocalization with Sec31A by IF. A validated clone was selected for use in experiments.

All primers and gene blocks used for cloning, Gibson assembly, and sequence confirmation were purchased from IDT. Confirmation of DNA sequences was done via Sanger sequencing by Genewiz.

### Primer and Gene Block Sequences

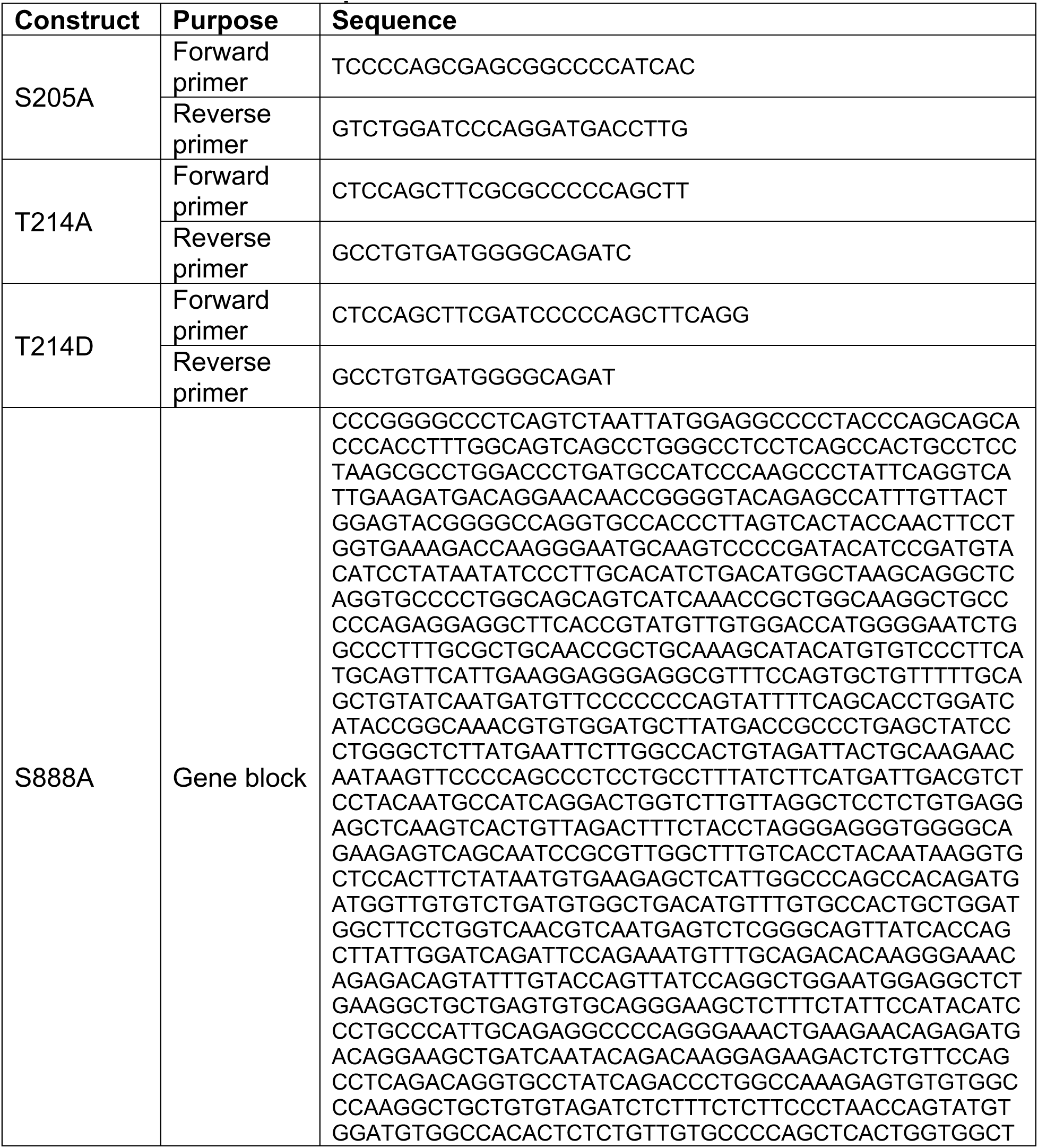
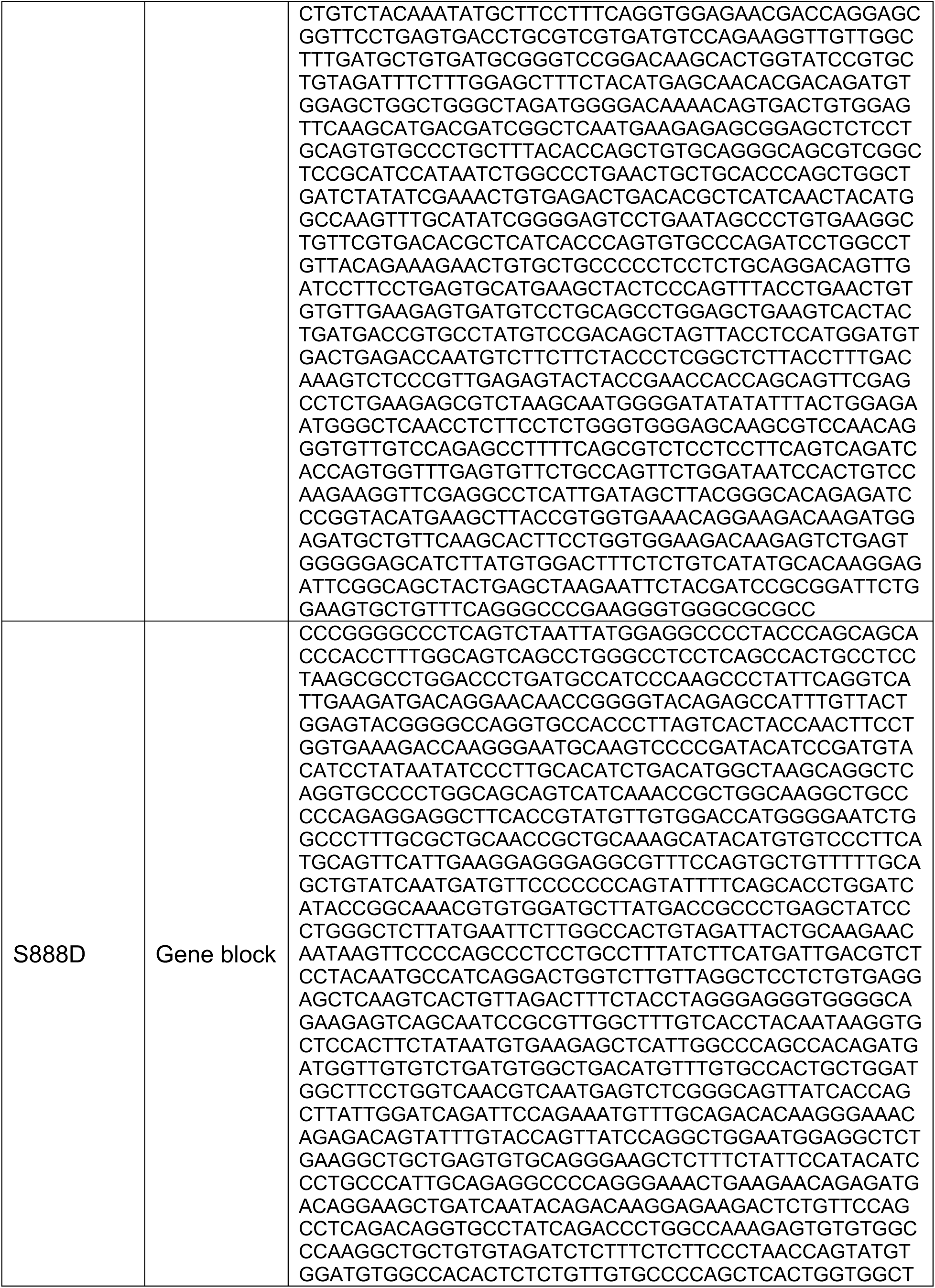
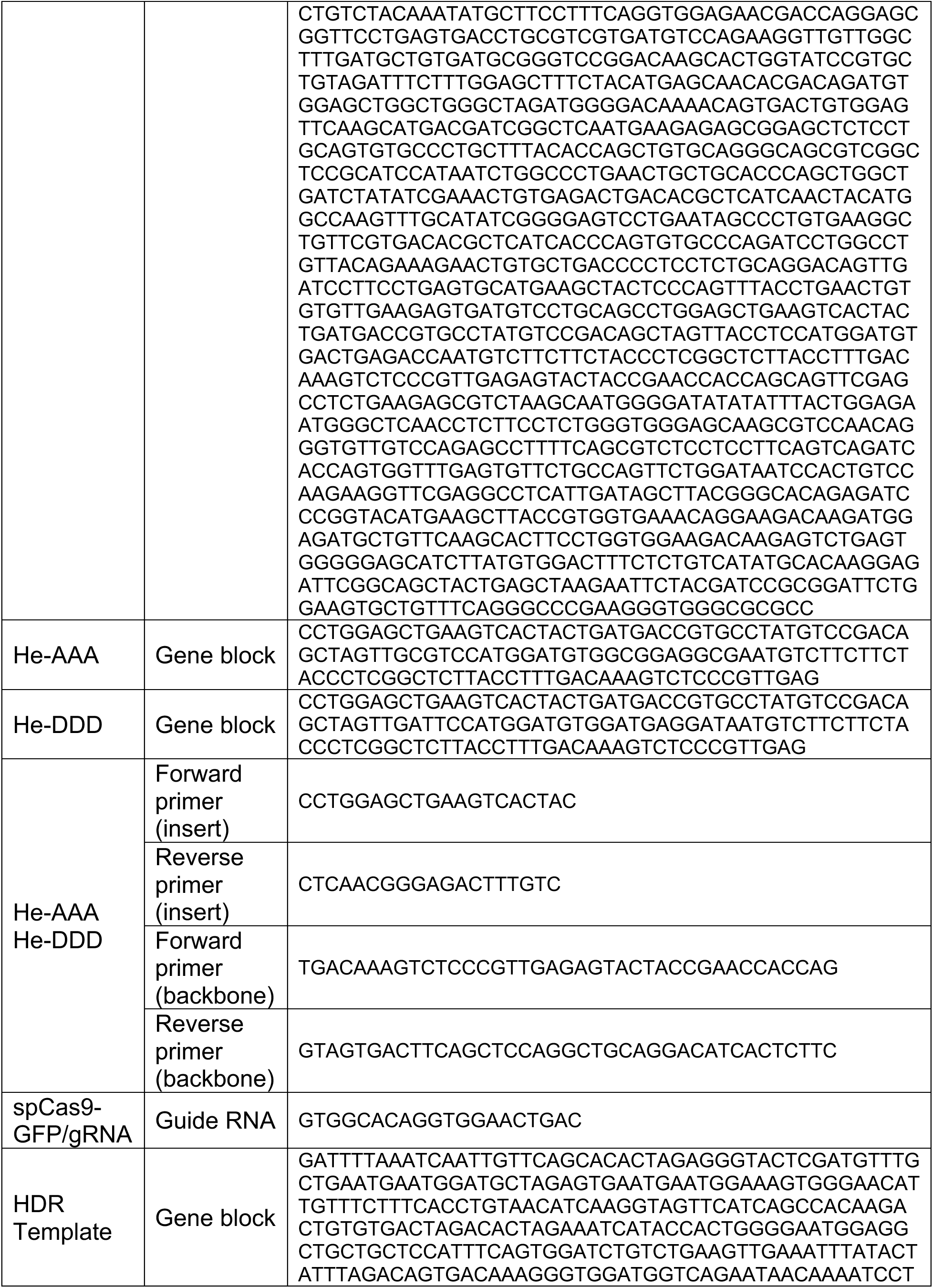
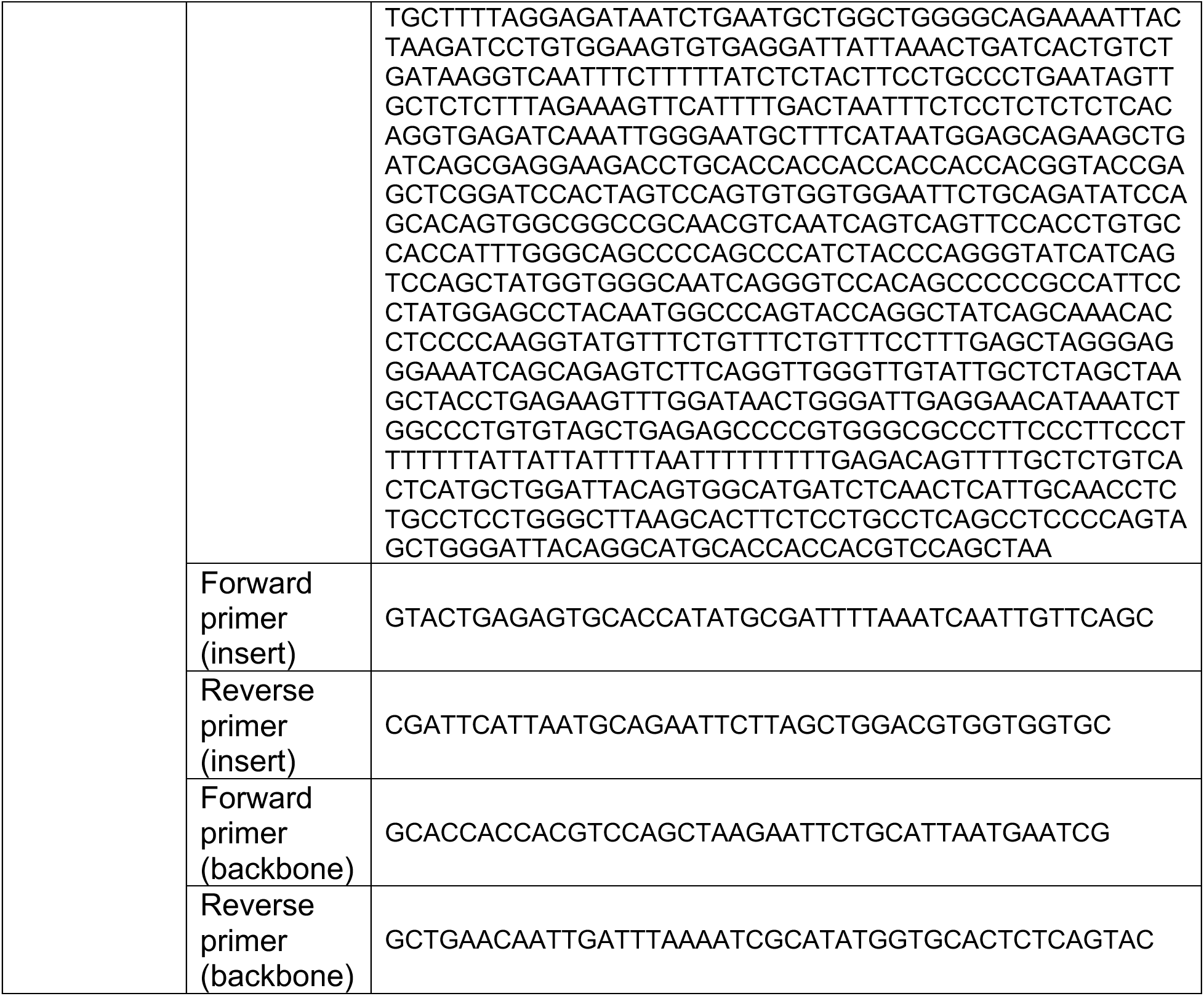

### Cell Culture and Cell Cycle Synchronization

Cells were maintained in Dulbecco’s modified Eagle Medium (DMEM, Sigma-Aldrich, D6429) containing 10% heat-inactivated fetal bovine serum (FBS, Sigma-Aldrich, F0926), 100 units/mL penicillin, and 100 μg/mL streptomycin (Pen/ Strep, Gibco, 15140-122) in a humidified incubator at 37 °C and 5% CO_2_.

Cells were synchronized by DTB as previously described **(60)**. Volumes of media, thymidine, and deoxycytidine were scaled depending on the size of culture plates.

### Nocodazole Arrest and OGA Inhibition

For nocodazole time range experiments, MH-Sec24C HeLa cells were grown to ∼80% confluence before addition of nocodazole to a final concentration of 3.3 µM or an equal volume of vehicle.

For compound-treated synchronized cells, 6.5 hours post-release, cells were treated with the OGA inhibitor, Thiamet-G, to a final concentration of 50 µM or an equal volume of vehicle (DMSO). Fifteen minutes later, cells were treated with nocodazole to a final concentration of 3.3 µM or an equal volume of vehicle. Untreated cells were harvested at 7 hours and 12.5 hours post-release to assess success of synchronization. Treated cells were harvested 10 hours post-release.

### Flow Cytometry

To assess DNA content and cell cycle profiles, cells were counted by hemacytometer. Then, ∼10^6^ cells per sample were spun down at 500 g for four minutes. Cell pellets were washed with 1x PBS, and spun again, after which pellets were resuspended in 300 µL 1x PBS and chilled on ice. To the chilled cells, 700 µL of ice-cold 100% ethanol (EtOH) was added dropwise, with careful mixing between drops. Cells were left in the 70% EtOH solution overnight at 4 _°_C. To remove EtOH, cells were spun down at 700 g for 3 minutes. Pellets were washed with 1x PBS and spun. Finally, the pellet was resuspended and incubated for 1 hr at room temperature in a staining solution of 0.05% TritonX-100, 50 µg/mL RNase A, and 20 µg/mL propidium iodide in 1x PBS. Prepared cell solutions were protected from light until ready for analysis by a BD FACSCanto II flow cytometer.

### Transfections

Generally, transient transfections with constructs listed above were performed per manufacturers’ instructions. For IB/IP experiments with mutants, Sec24C knock-out (KO) HeLa cells **(34)** grown on 15 cm plates were transfected (Lipofectamine3000) at ∼60% confluence with 15 µg DNA per plate. Medium was aspirated and replaced with fresh, complete medium 12 hrs after transfection. For cycloheximide chase, cells in 6-well plates were transfected (TransIT-HeLaMONSTER) at 25% confluence with 2 µg DNA per well. 24 hrs after transfection, medium was aspirated, cells were washed twice with 1x PBS, and fresh, complete medium was added to wells. For IF experiments with mutants, cells in 12-well plates were transfected (TransIT-HeLaMONSTER) at 20% confluence with 1 µg DNA per well.

### Immunoblotting (IB) and Immunoprecipitation (IP)

IBs and IPs were performed as previously described **(33, 34)**, with the following changes. Lysates were prepared at final concentrations between 1-3 mg total protein per mL of lysis buffer. IP lysis buffer containing SDS was not used in lysate preparations. However, phosphatase inhibitor cocktail was included in most lysis buffers. For all myc IPs, the mouse anti-myc (9E10) antibody was used, and 2-4 µg myc antibody was used per mg of total protein. Mock IPs were performed using equal quantities of mouse anti-FLAG antibody in place of mouse anti-myc (9E10). After IPs, beads were washed 5x with IP lysis buffer without EDTA and without phosphatase inhibitors to allow downstream compatibility with Phos-Tag SDS-PAGE. IP lysis buffer without EDTA was also used to dilute 5x loading dye to 1x for eluting proteins off beads. Rabbit anti-myc (71D10) was used to blot for myc unless otherwise noted. In addition to washes with 1x Tris-buffered saline with Tween-20 (TBST), nitrocellulose membranes were washed three times, 5 minutes each wash, in 1x TBS, directly prior to image acquisition by LiCor Odyssey CLx Imager.

Additionally, “dual-blotting” was employed, where useful, to blot for myc and O-GlcNAc simultaneously. Primary antibodies for myc (rabbit) and either 18B10 (mouse) or 9D1 (mouse) were combined and exposed to membranes simultaneously. Two secondary antibodies with anti-rabbit (anti-rabbit IgG IRDye 680RD) or mouse IgG (anti-mouse IgG IRDye 800CW) specificity, and each conjugated to a unique fluorophore, were then used to distinguish myc from O-GlcNAc signals on a single blot. Dual-blotting did not necessitate any changes to the procedure more generally and produced identical results to single-antibody blots when compared.

### IB Quantification

For quantification of total protein in cell lysates, O-GlcNAc levels on IP-ed Sec24C, or co-IP of Sec23A/B with Sec24C, images acquired with the LiCor Odyssey CLx Imager were exported as TIFF files and analyzed using Fiji/ImageJ. Background subtraction was applied using a rolling ball radius equal to the length of the widest band-of-interest within the image. Despeckling was also applied. Lanes were highlighted, and integrated band intensities were quantified using Fiji’s Gel Analyzer tool. For total protein quantification, quantified protein signal was divided by tubulin or GAPDH signal from the same sample (except for the cycloheximide chase experiment). For O-GlcNAc and co-IP quantification, the same process was followed, but quantified O-GlcNAc and Sec23A/B signals, respectively, were instead divided by quantified Sec24C signal from the same sample.

### Phos-Tag SDS-PAGE and On-Bead Phosphatase Assay

Mn^2+^ Phos-Tag gels were prepared in-house according to manufacturer’s instructions (Fujifilm Wako Pure Chemical Corporation). Gel electrophoresis was conducted as soon as possible after casting and allowed to run overnight at 35 V. To confirm that slower-migrating bands were the result of phosphorylation, Sec24C captured on washed Protein A/G-beads was incubated with λ phosphatase for 45 min at 30 _°_C. The bead slurry was occasionally mixed by gentle vortexing during the incubation period. After incubation, 5x loading dye was added to samples to a final concentration of 1x, and samples were heated at 95 _°_C for five minutes.

### Cycloheximide Chase

Following transfection with either WT, He-AAA or He-DDD Sec24C as described above, cells were treated with cycloheximide to a final concentration of 50 µg/mL for 0, 1, 2, 4, 8, 20 or 24 hours before harvest.

### Endogenous Sec24C Purification for PTM Mapping

Tandem myc IP and nickel-NTA column purification was performed on DTB-synchronized MH-Sec24C HeLa cells as previously described **(34, 81)**. 20 to 25 15-cm plates were used per condition tested, with conditions being 4 hrs (S phase), 7 hrs (G2), 9 hrs (mitosis), and 12 hrs (G1) post-release. 50 to 125 mg total protein was harvested per condition per replicate for MH-Sec24C purification.

### LC-MS/MS

Eluted samples in PBS were supplemented with 5% SDS, reduced for 15 min at 80 _°_C with 10 mM dithiothreitol, and subjected to SDS-PAGE separation on a 4-12% bis-tris NuPAGE gel (Invitrogen). Bands corresponding to Sec24C were excised and subjected to in-gel tryptic digestion using standardized protocols **(103)**. Digested peptides were lyophilized to dryness and resuspended in 12 µL of 0.2% formic acid/2% acetonitrile. Each sample was subjected to chromatographic separation on a Waters MClass UPLC equipped with a 1.7 µm HSS T3 C_18_ 75 µm I.D. X 250 mm reversed-phase column. The mobile phase consisted of (A) 0.1% formic acid in water and (B) 0.1% formic acid in acetonitrile. 3 µL was injected and peptides were trapped for 3 min on a 5 µm Symmetry C_18_ 180 µm I.D. X 20 mm column at 5 µl/min in 99.9% A. The analytical column was then switched in-line and a linear elution gradient of 5% B to 40% B was performed over 90 min at 400 nL/min. Data collection on the Fusion Lumos mass spectrometer operating in a data-dependent acquisition (DDA) mode of acquisition with a r=120,000 (@ m/z 400) full MS scan from m/z 350 – 1800 with a target AGC value of 4e5 ions was performed. MS/MS scans with HCD settings of 32% were acquired in Orbitrap at r=15,000 (@ m/z 400) with a target AGC value of 1e4 and max fill time of 105 ms. If HexNAc losses (204.0867, 138.0545, or 366.1396) were found in an MS/MS spectrum, an ETD activated scan was triggered with AGC settings of 4e5 ions with accumulation of 120ms. A 20s dynamic exclusion was employed to increase depth of coverage.

Raw LC-MS/MS data files were processed in Proteome Discoverer 3.0 (ThermoFisher) and then submitted to independent Sequest database searches against a human protein database containing both forward (20,260 entries) and reverse entries of each protein. Search tolerances were 2 ppm for precursor ions and 0.02 Da for product ions using trypsin specificity with up to two missed cleavages. Dynamic mass modifications on S/T corresponding to GlcNAc (204.0867m/z) were included. All searched spectra were imported into Scaffold (v5.3, Proteome Software) and scoring thresholds were set to achieve a peptide false discovery rate of 1% using the PeptideProphet algorithm. Raw data were also imported into Protein Discoverer 3.0 for area under the curve measurements of eluting peptides. Extracted ion chromatograms of individually GlcNAcylated peptides were visualized within Skyline (MacCoss Laboratory, Univ of Washington).

### Sequence Alignments and Phylogeny

For determination of amino acid conservation, sequences of Sec24C homologs in the NCBI database were selected from a broad array of animal phyla, with some fungal and plant homologs included. Multiple sequence alignment (MSA) was conducted using Geneious Prime’s aligner tool, which employs a progressive pairwise alignment algorithm. The tool’s default parameters were used (global alignment with free end gaps and a Blosum62 cost matrix). Aligned sequences were manually assessed for conservation of either serine or threonine residues at identified PTM sites in human Sec24C. Amino acid conservation between Sec24 human paralogs was determined the same way. From the MSA, a phylogenetic tree was prepared with Geneious Prime’s tree builder tool using a Jukes-Cantor genetic distance model and neighbor-joining with no outgroup included.

### IF and Image Acquisition

Cells were seeded and grown on circular cover glass (#1.5, 18 mm) within 12-well plates. If required, transfections were performed as described above. Cells were fixed and permeabilized by gentle pipetting of 100% methanol to cover slips, followed by a 20-minute incubation at 4 _°_C. After this, methanol was removed, and cover slips were covered with cold 1x PBS and washed for five minutes with gentle agitation. This wash was repeated a total of four times. The slips were then incubated for 1 hr at room temperature with rotation in blocking buffer (5% goat serum, 0.3% Triton X-100, and 0.05% NaN_3_ in 1x PBS). Directly after blocking, slips were incubated overnight at 4 _°_C with primary antibodies diluted in blocking buffer (dilutions for antibodies used are detailed in the chart above). The following day, slips were washed as before, after which they were incubated for two hours in the dark at room temperature with secondary antibodies diluted in blocking buffer (dilutions detailed above). Slips were then washed four times with 1x PBST (1x PBS + 0.1%Triton X-100). Finally, slips were affixed to slides with mounting media containing DAPI and allowed to cure in the dark overnight before being imaged the following day.

Imaging was conducted on a Zeiss 710 inverted confocal microscope using either a 40x (1.30 Oil EC Plan Neofluar DIC) or 63x oil immersion (1.40 oil Plan-Apochromat DIC) objective lens. Diode (405 nm), argon ion (488 nm), and helium-neon (594 nm) lasers were used for fluorophore excitation. 1912×1912 pixel resolution images were acquired by frame-scanning bidirectionally using the imaging mode in Zeiss Zen Black Acquisition software. Z-stacks were taken with a step size of 0.36 µm for mutant-related experiments, and a step-size of 0.6 µm for cell cycle-related experiments, in both cases with 4 images taken across the Z-plane. Detection ranges for fluorophores were determined to maximize detection and minimize cross-talk with the assistance of ThermoFisher Fluorescence SpectraViewer (https://www.thermofisher.com/order/fluorescence-spectraviewer/#!/).

### Image and Particle Analysis

IF image analysis was performed on Fiji/ImageJ for cell cycle-related experiments, and on Imaris for mutant-related experiments. Imaris’s “Surface” feature was used to define Sec24C and Sec16A regions. Surfaces smaller than ∼200 nm in diameter were excluded from further analysis. The volumes of remaining Sec24C surfaces, as well as their distance to the nearest neighboring Sec16A surface were calculated. Additionally, Sec24C surfaces were tallied per image, and the ratio of surfaces overlapping Sec16A to total Sec24C surfaces was calculated with the aid of JMP Pro 17.

For quantification with Fiji/ImageJ, MAX Z-projection was applied to flatten 3D images. Regions of interest (ROI) were manually drawn to select for individual cells. ROIs were then subjected to automatic thresholding (Otsu), and Sec24C attributes were determined using the “Measure” and “Analyze Particles” features of Fiji/ImageJ as needed. An ImageJ Macro was developed and utilized to streamline analysis (see Data Availability for access to code).

For quantification of O-GlcNAc signal in Figure S3, after applying a MinError threshold, total RL2 intensity was measured across an entire image, and divided by the total area of cells within the image. A unique Jython (ImageJ Python) script was developed with the assistance of the open access version of OpenAI’s ChatGPT-4 to streamline this process (see Data Availability for access to code).

### Statistical Analysis

All statistical analyses were performed using GraphPad Prism version 10, with details of statistical tests described in figure legends. All data represent a minimum of three independent biological replicates.

## Data Availability

Sec24C O-GlcNAc site-mapping data have been uploaded to the ProteomeXchange Consortium using the PRIDE partner repository with accession number PXD060692.

All Python code used for image analysis has been uploaded for public access to GitHub and has been assigned the following DOI via Zenodo: 10.5281/zenodo.14861391.

## Supporting information

This article contains supporting information.

## Acknowledgments

HeLa cells, established through the non-consensual harvesting of cervical cancer cells from Henrietta Lacks, a Black American woman, were seminal to the work of this paper as well as biomedical research in general. As such, we would first like to acknowledge and pay solemn tribute to the contributions of Henrietta Lacks, now deceased, and the Lacks family. We would also like to thank Dr. Mike Cook, now deceased, and Dr. Ying Guo for help with cell sorting and flow cytometry; Professor Ted Slotkin and Jimin Hu for insights into statistical analysis; Dr. So Young Kim for the cloning of Cas9/gRNA constructs; and Drs. William Kasberg and Ben Carlson for assistance with immunofluorescence techniques and analysis. We acknowledge the use of OpenAI’s ChatGPT-4 by G.R.G. to generate Python code for image analysis by ImageJ (see Data Availability section for code access information).

## Funding and additional information

This research was supported by funds from the National Institutes of Health grant R01GM117473 (to M.B.). The content is solely the responsibility of the authors and does not necessarily represent the official views of the National Institutes of Health.

## Conflicts of interest

The authors declare that they have no conflicts of interest with the contents of this article.

## Abbreviations

The abbreviations used are

CDK1: cyclin-dependent kinase 1
CK1δ: casein kinase 1 isoform delta
COPII: coat protein complex 2
cTAGE5: cutaneous T-cell lymphoma-associated antigen 5
DMSO: dimethyl sulfoxide
DTB: double thymidine block
ERES: endoplasmic reticulum exit sites
HRP: horseradish peroxidase
IF: immunofluorescence
IP: immunoprecipitation
LC-MS/MS: liquid chromatography tandem mass spectrometry
Ni-NTA: nickel-nitrilotriacetic
O-GlcNAc: O-linked β-*N*-acetylglucosamine
pH3: phospho-histone-H3
PLK1: polo-like kinase 1
PP1/PP2A: protein phosphatase 1/2A
PTM: post-translational modification
TANGO1: transport and Golgi organization protein 1
TG: Thiamet-G.

## Supporting Information

**Figure S1.**
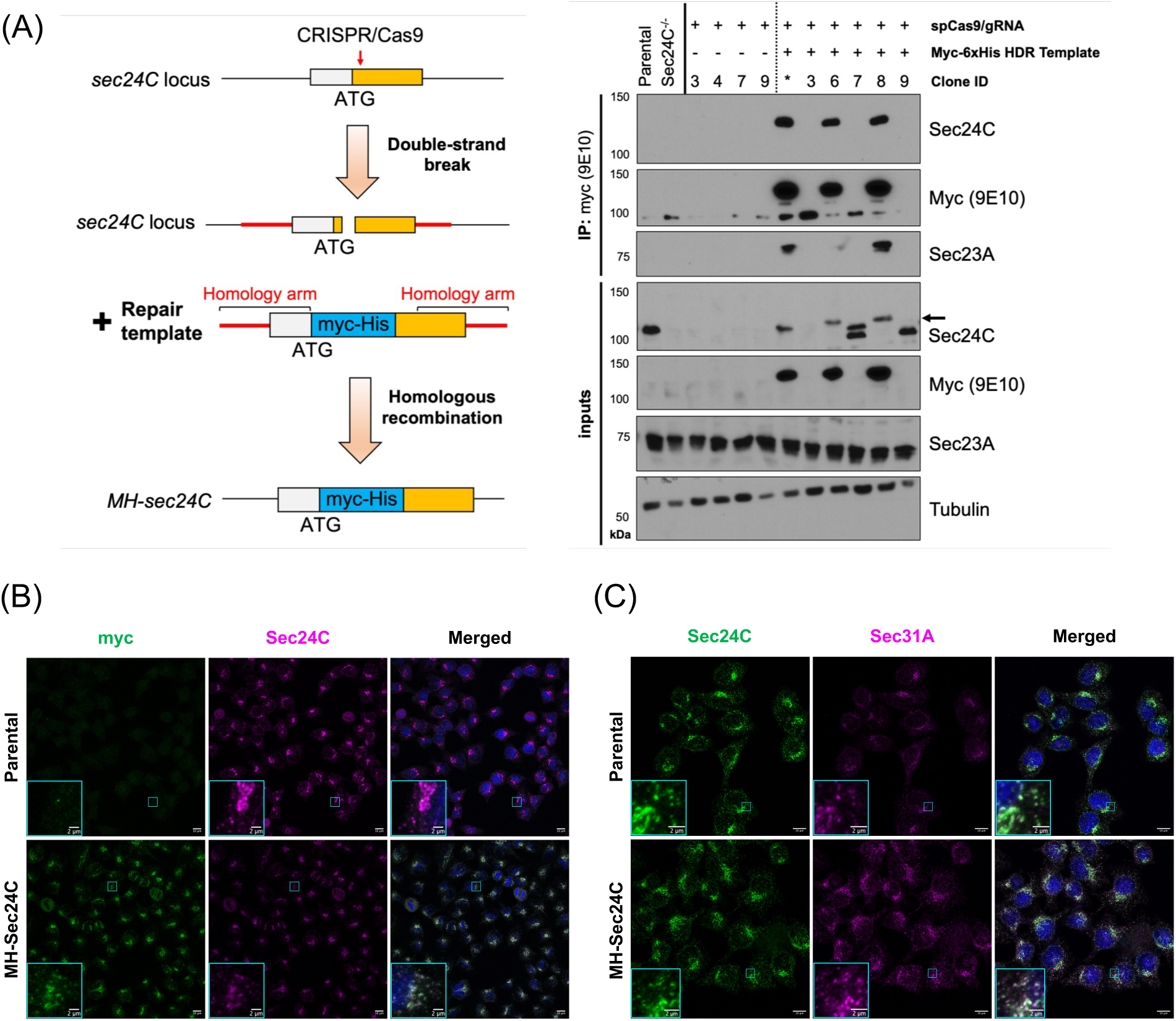
(A) CRISPR-Cas9 technology was used to add a myc-6xHis tag to the N-terminus of Sec24C at its endogenous locus in HeLa cells (left). Myc-6xHis-tagged single-cell derived clones were identified and confirmed by IP/IB (right). Cells were transfected with a plasmid encoding GFP-tagged sp-Cas9 and a guide RNA (gRNA) sequence targeting the N-terminus of Sec24C, plus a repair template encoding myc-6xHis with homology arms flanking the spCas9 cut-site (myc-6xHis homology directed repair [HDR] template). GFP-positive single cells transfected with spCas9-GFP alone or spCas9-GFP and HDR template were sorted by flow cytometer. Cells were cultured until sufficient for IP/IB. Clone IDs represent unique single-cell-derived colonies. A transfected, unsorted mixed population was included as a control (denoted *). Clone 8 was selected for further studies and was renamed MH-Sec24C. Black arrow indicates correctly tagged Sec24C. (B, C) Colocalization of myc/Sec24C (B), and Sec24C/Sec31A (C) in MH-Sec24C and parental HeLa cells, as assessed by IF. Scale bar = 10 µm, inset scale bar = 2 µm.

**Figure S2.**
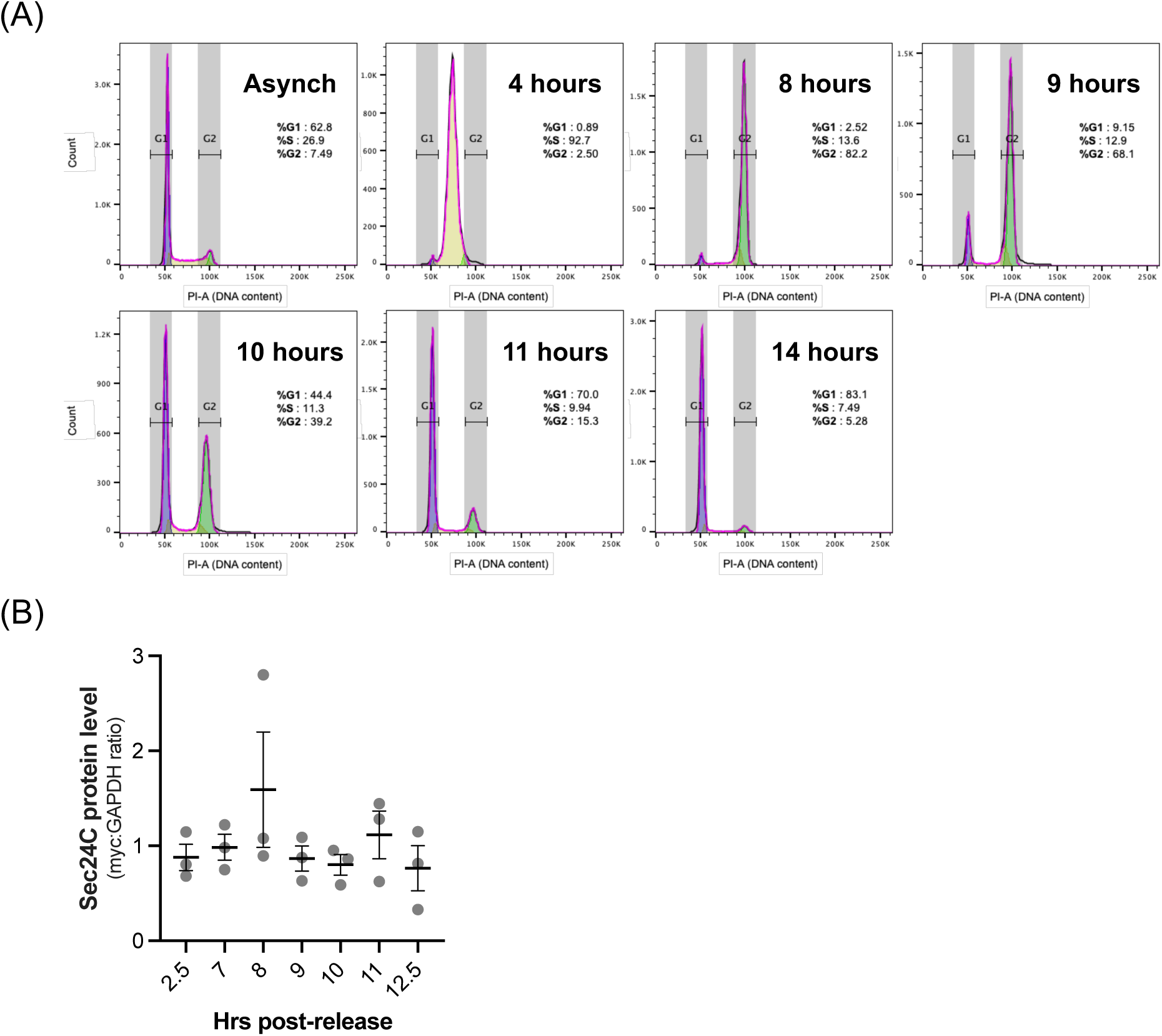
(A) MH-Sec24C cells were synchronized by DTB, and cell cycle profile was assessed by propidium iodide (PI) staining of DNA content and flow cytometry at the indicated times post-DTB release. (B) Levels of myc-6xHis-Sec24C were quantified, normalized to GAPDH, and compared across cell cycle stages by one-way ANOVA with Tukey’s post-hoc test. *p* values > 0.05 are not displayed.

**Figure S3.**
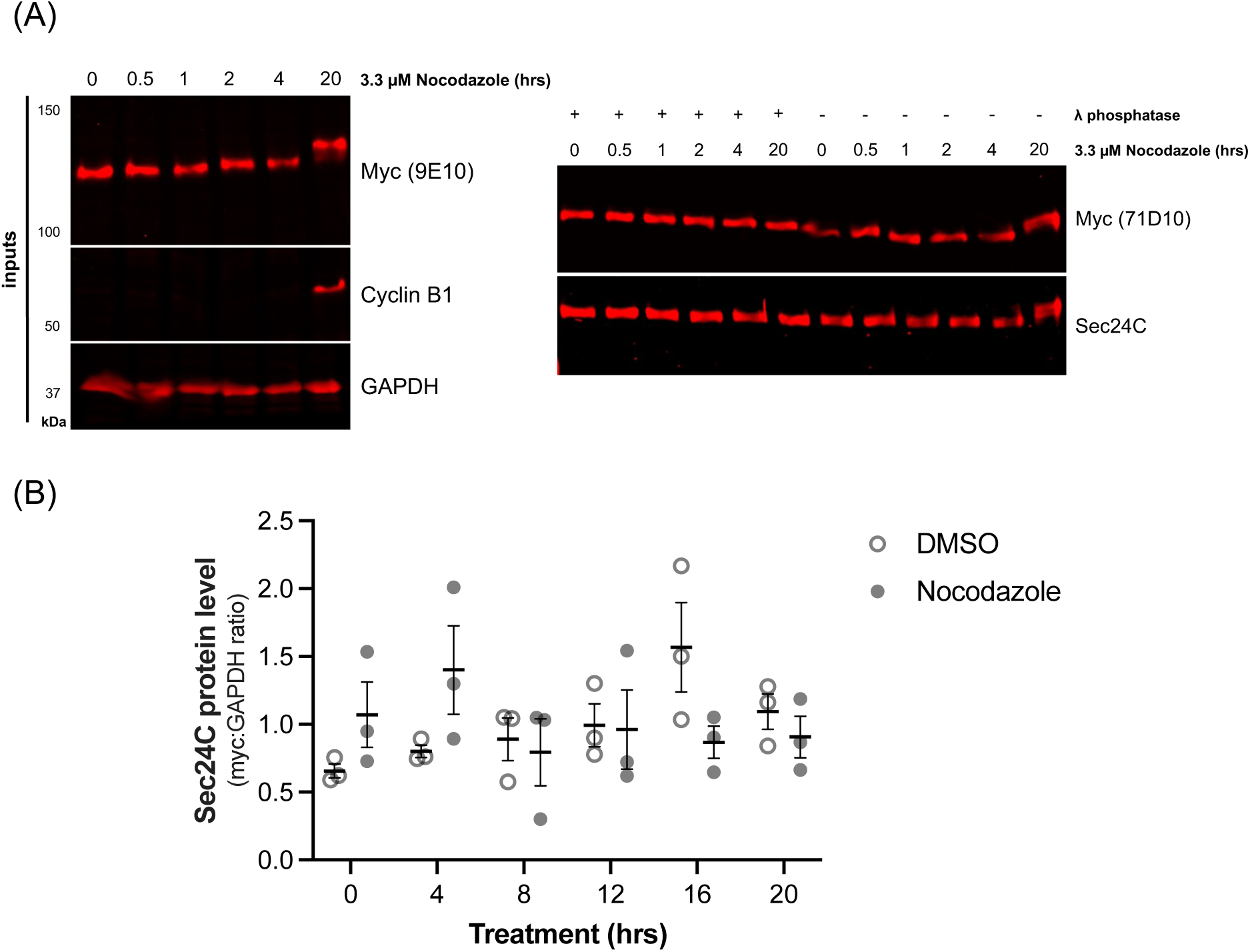
(A) MH-Sec24C cells were treated with 3.3 µM nocodazole for the indicated times, and lysates were treated with λ phosphatase (or mock conditions) and were analyzed by myc IP and IB. (B) Levels of myc-6xHis-Sec24C from (A), plus corresponding DMSO control samples, were quantified, normalized to GAPDH, and analyzed by two-way ANOVA with Tukey’s post-hoc test. *p* values > 0.05 are not displayed.

**Figure S4.**
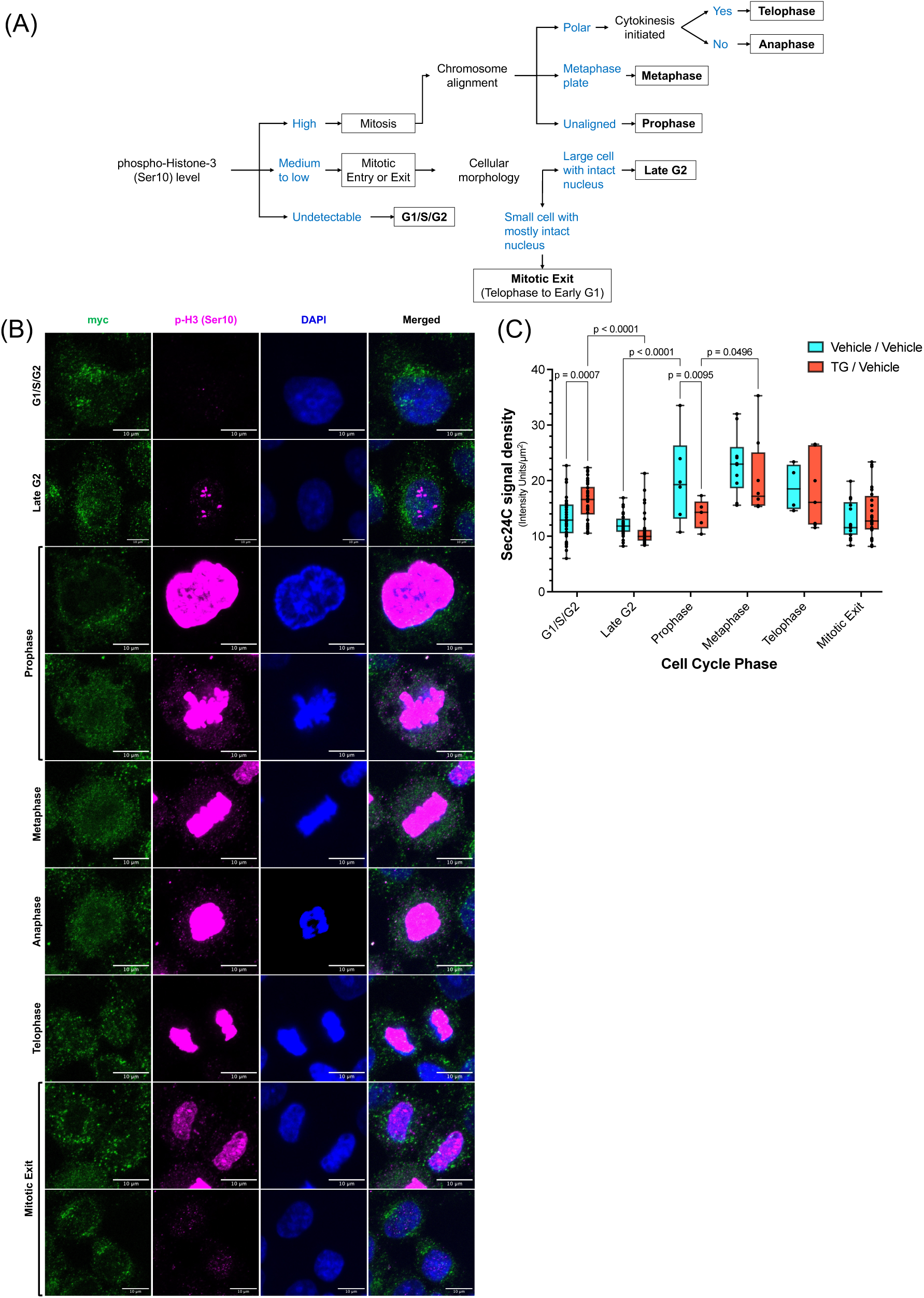
(A) Flow-chart describing classification of imaged cells into cell cycle phase groups by phospho-histone-H3 (Ser10) staining and nuclear morphology. (B) Representative IF images of MH-Sec24C cells in each identifiable cell cycle phase. Scale bars = 10 µm. (C) Total intensity of Sec24C particles within a cell divided by area of the cell. Data are presented with medians indicated by solid lines, 25th-75th percentile range by boxes, and full range from minimum to maximum by whiskers. Anaphase was excluded from the analysis due to insufficient numbers of anaphase cells. Statistical significance was assessed by two-way ANOVA followed by Tukey’s post-hoc test. Only comparisons between TG and Vehicle and consecutive cell cycle phases are shown. *p* values > 0.05 are not displayed.

**Figure S5.**
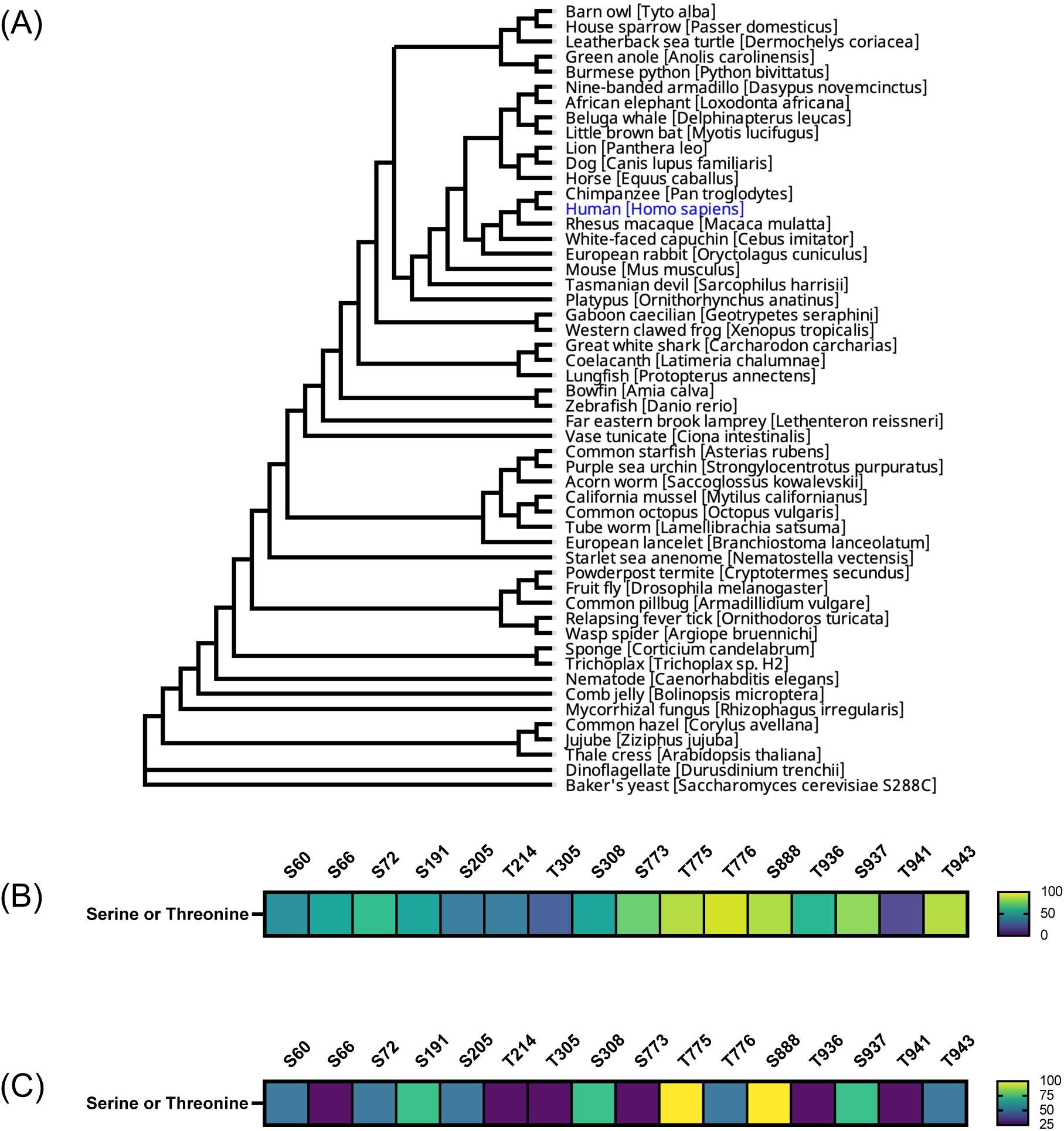
(A) Phylogenetic tree of Sec24C orthologs selected from a wide array of eukaryotic clades. Each ortholog analyzed represents the protein within that species that is most similar to the human Sec24C amino acid sequence. (B) Heatmap illustrating the evolutionary conservation of serine/threonine residues at the cognate positions of human Sec24C PTM sites in the Sec24C orthologs listed in (A). (C) Heatmap illustrating the conservation of serine/threonine residues at the cognate positions of identified and candidate Sec24C PTM sites in the four human Sec24 paralogs. Color scales in (B) and (C) indicate the percentage of serine/threonine conservation at these sites, as determined by the alignments.

**Figure S6.**
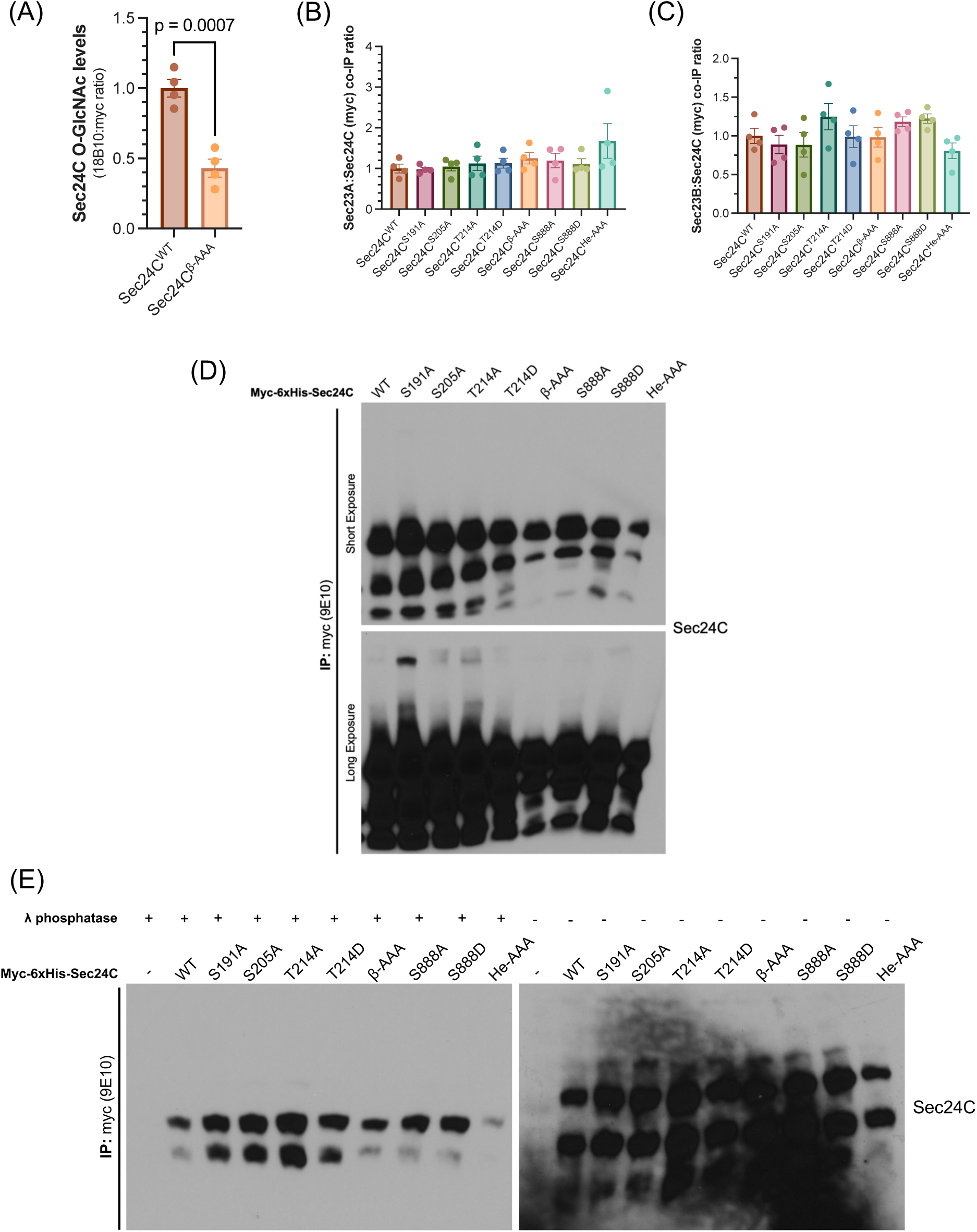
(A) Data for O-GlcNAc levels of WT and β-AAA Sec24C from the experiment depicted in Figure 7A were analyzed by Student’s t-test. Data were normalized to the WT mean for each replicate (n=4). (B, C) Levels of Sec23A (B) and Sec23B (C) that co-IP-ed with Sec24C in the experiment depicted in Figure 7A were quantified. Data points represent independent biological replicates (n=3 or 4) normalized to the WT mean for each replicate. Bars represent means ± SEM. Statistical significance was assessed by one-way ANOVA, and post-hoc comparisons were conducted between selected groups (WT versus all other constructs and Ala versus Asp mutants of the same residue) using Šídák’s method. *p* > 0.05 comparisons are not shown. (D) Sec24C KO cells were transfected as indicated, and lysates were analyzed by myc IP, Phos-Tag SDS-PAGE and IB. (E) Cell lysates were prepared as in (D), treated with λ phosphatase or control conditions, and analyzed by Phos-Tag SDS-PAGE and IB. Lysates used in (D) and (E) are identical to those in Figure 7A.

**Figure S7.**
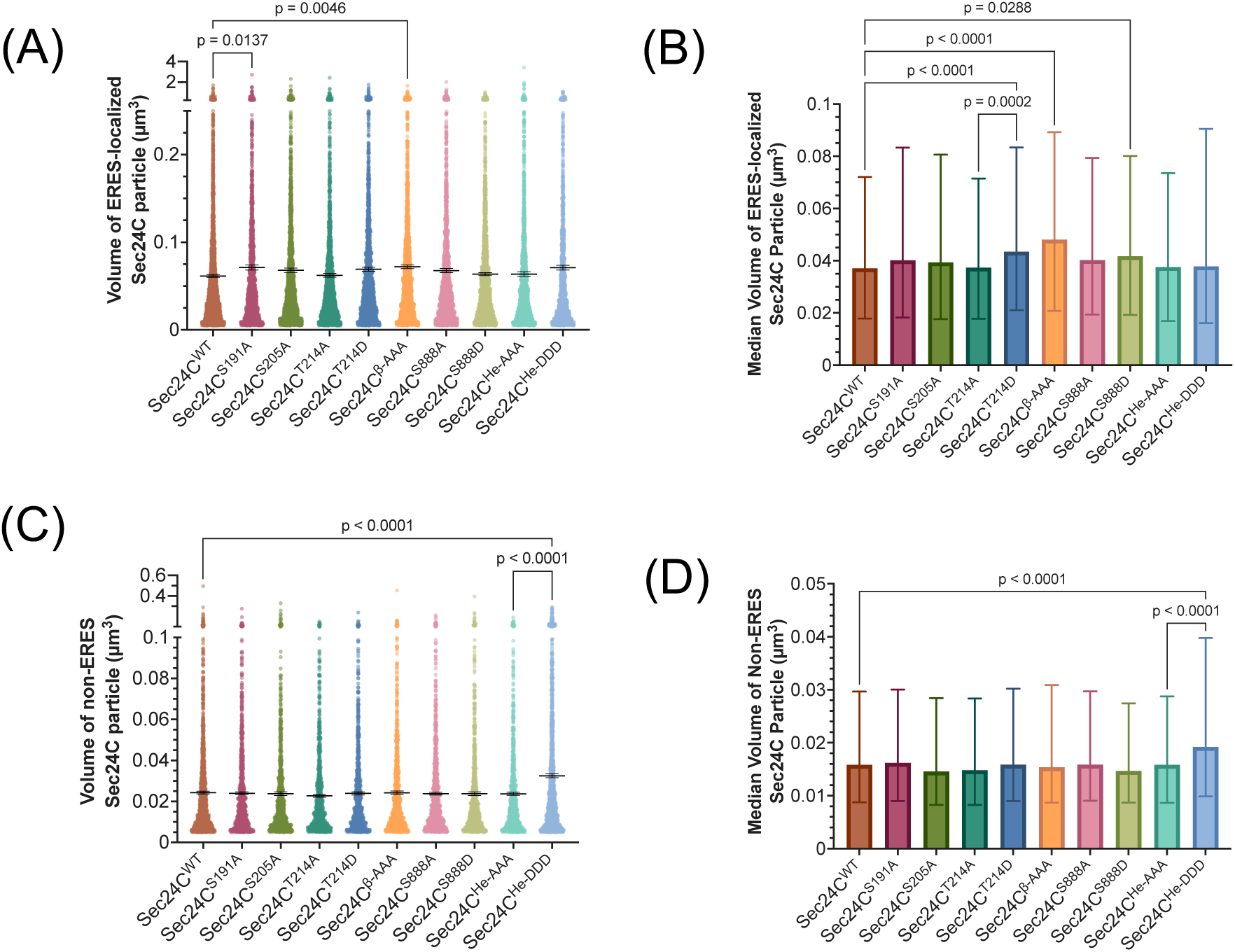
From the IF data depicted in Figure 9A, (A, C) mean and (B, D) median volumes of individual Sec24C particles were calculated, relative to Sec16A-marked ERES. Means are displayed as black lines ± SEM within scatter plots of all values, and medians are displayed as bars ± interquartile ranges. For each replicate (n=3 to 5) and Sec24C construct tested, 998 Sec24C particles were measured. Differences between means were assessed by one-way ANOVA and Šídák’s multiple comparisons test. Differences between medians were assessed by Kruskal-Wallis test, followed by Dunn’s multiple comparison. Comparisons were only computed for selected combinations (WT versus all other constructs and Ala versus Asp mutants of the same residue). *p* > 0.05 comparisons are not shown.

## Notes

### Competing Interest Statement

The authors have declared no competing interest.

